# Aged blood inhibits hippocampal neurogenesis and activates microglia through VCAM1 at the blood-brain barrier

**DOI:** 10.1101/242198

**Authors:** Hanadie Yousef, Cathrin J Czupalla, Davis Lee, Ashley Burke, Michelle Chen, Judith Zandstra, Elisabeth Berber, Benoit Lehallier, Vidhu Mathur, Ramesh V Nair, Liana Bonanno, Taylor Merkel, Markus Schwaninger, Stephen Quake, Eugene C Butcher, Tony Wyss-Coray

## Abstract

An aged circulatory environment can promote brain dysfunction and we hypothesized that the blood-brain barrier (BBB) mediates at least some of these effects. We observe brain endothelial cells (BECs) in the aged mouse hippocampus express an inflammatory transcriptional profile with focal upregulation of Vascular Cell Adhesion Molecule 1 (VCAM1), a protein that facilitates vascular-immune cell interactions. Concomitantly, the shed, soluble form of VCAM1 is prominently increased in the aged circulation of humans and mice, and aged plasma is sufficient to increase VCAM1 expression in cultured BECs and young mouse hippocampi. Systemic anti-VCAM1 antibody or genetic ablation of VCAM1 in BECs counteracts the detrimental effects of aged plasma on young brains and reverses aging aspects in old mouse brains. Thus, VCAM1 is a negative regulator of adult neurogenesis and inducer of microglial reactivity, establishing VCAM1 and the luminal side of the BBB as possible targets to treat age-related neurodegeneration.

---

Brain structure and function deteriorate with age, steadily driving cognitive impairments and susceptibility to neurodegenerative disorders in humans ^1^. How aging leads to these manifestations is poorly understood but two cellular hallmarks of brain aging are particularly noticeable. The first is an increase in activation of innate immune pathways and an increase in the activation state of microglia and astrocytes, frequently referred to as “neuroinflammation” ^2–4^. The second aging hallmark is the precipitous loss of stem cell numbers and activity in the dentate gyrus (DG) of the hippocampus, one of two neurogenic regions of the adult mammalian brain ^5^. The hippocampus is crucial for learning and memory, and is particularly vulnerable to age-related neurodegeneration and diseases such as Alzheimer’s disease (AD) ^6^. Hippocampal neurogenesis is important for some aspects of learning and memory, especially as it relates to spatial navigation and pattern separation ^7^.

While many of these age-related changes in the brain may be the consequences of cell-intrinsic and brain-localized mechanisms of aging, we asked if changes in secreted signaling proteins, dubbed the communicome ^8^, could be used to understand, characterize, and quantify aspects of brain aging and cognitive impairment. Indeed, such changes in plasma or CSF proteomes are not only abundant with aging and disease ^9,10^, but factors in young blood or plasma from mice or humans are sufficient to increase brain function in the hippocampus ^9,11,12^ and the subventricular zone (SVZ) ^13^. Conversely, mice exposed to old blood showed reduced neurogenesis in the hippocampus ^9,14^. At least some of the inhibitory effects can be attributed to the chemokine CCL11 ^9^ and the MHC class I associated molecule β2-microglobulin (B2M; ^15^), consistent with a general increase in inflammatory mediators with aging ^16^. Considering the tight regulation of transport of molecules across the BBB and its role as a protective barrier with limited permeability to macromolecules ^17^, it is currently unclear how pro-youthful or pro-aging factors may modulate brain function ^1^. Here, we describe a mechanism by which VCAM1 expressed on the luminal (blood-facing) side of the BBB is necessary to mediate the inhibitory effects of aged plasma on hippocampal neurogenesis in mice and humans and the pro-inflammatory effects of aged plasma on microglia. Furthermore, systemic administration of anti-VCAM1 antibody prevented the adverse effects of aged plasma and rejuvenated aged mice brains. We propose that VCAM1 is a potential therapeutic target of age-related neurodegeneration.

## Results

### Aged BECs are transcriptionally activated

To determine the transcriptional changes associated with aged BECs, we acutely isolated primary CD31+ BECs from young and aged pooled mouse cortices and hippocampi and analyzed their transcriptome using RNA sequencing **(Supplementary Fig. 1a-b**). Unsupervised cluster analysis revealed prominent age-dependent changes in the transcriptome with over 1000 differentially expressed genes (**Fig. 1a**). Cell purity was confirmed based on high gene expression values for BEC specific genes, and very low or undetectable expression of other CNS cell type-specific markers (**Fig. 1b**, **Supplemental Fig. 1c**). GeneAnalytics Pathway Analysis of differentially expressed genes revealed numerous pathways associated with aging (**Supplemental Table 1**), including gene pathways associated with cell adhesion, immune cell activation, stress response and vascular remodeling ^18–21^ linking them to inflammation, leukocyte adhesion, BBB maintanence and inhibition of cell proliferation. Analysis of the highly expressed and differentially expressed transcripts revealed an inflammatory and activated profile with age as illustrated by the doubling in mRNA expression of the MHC class I molecules β2-microglobulin *(B2m)* and *H2-K1*, two of the most highly expressed transcripts in BECs (**Fig. 1c**). *Tspo*, a marker of neuroinflammation commonly used in human PET scans to assess the level of microglial activation in neurodegenerative diseases ^22^, was highly expressed in BECs and also significantly increases with age, as was *Vwf*, a blood glycoprotein involved in hemostasis, elevated under acute and chronic inflammation and known to promote vascular inflammation ^23^ (**Fig. 1c**). This suggests that aged BECs are in an activated, inflammatory state.

**Figure 1.**
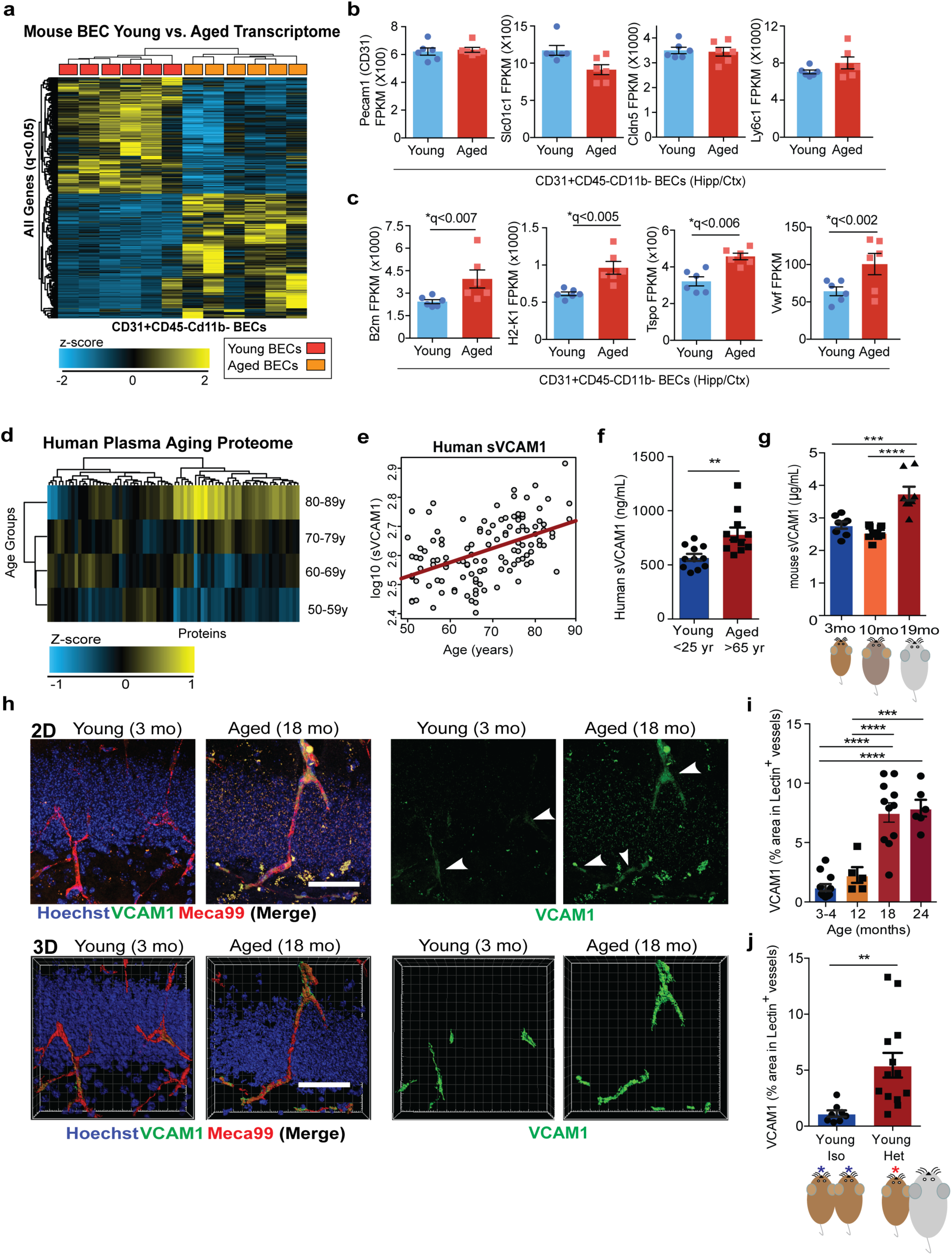
BECs are activated with age. Systemic and cerebrovascular VCAM1 increases with aging and heterochronic parabiosis. (a) Heat map displaying up or down-differentially regulated genes in young versus aged BECs. There were 1006 significant differentially expressed genes (*q<0.05). (b) Fragments Per Kilobase of transcript per Million mapped reads (FPKM) of BEC cell-type specific markers. (c) FPKM values of inflammation and activation related genes. (d) Heat map showing changes in 31 out of 74 human plasma factors with aging (p<0.05, Spearman’s correlation coefficient). Multiplex assay used (n=118 healthy humans). VCAM1 was the top factor (spearman’s correlation coefficient=0.47, p=7.7e-08). (e) Spearman correlation of VCAM1 levels and age with a significant inflection point around age 65 (Spearman’s correlation coefficient = 0.047; q< 6×10^-6^). (f) Human sVCAM1 ELISAs in 11 young (<25 years old) or 11 aged (>65 years old) plasma from healthy donors. **p<0.005, Student’s *t-test*. (g) ELISA for mouse sVCAM1 in plasma from young (3-month-old; n=8), middle-aged (8–10-month-old; n=10), and aged (18-month-old; n=8) mice. ****p<0.0001, 1-way ANOVA. (h) Representative confocal images in the DG of young (3-month-old) or aged (18-month-old) mice given retro-orbital (r.o.) injections of fluorescently conjugated anti-VCAM1 and anti-Meca99 2 hours before perfusion. Hoechst labels cell nuclei. Scale bar = 50 μm. 3D rendering of the 2D images are displayed. 3D Scale bar = 50 μm. (i) Quantification of VCAM1+Lectin+ stained brain vasculature in young, middle, and aged hippocampi. n=12 young (3–4-month-old), 6 aged (18-month-old), and 6 very aged (24-month-old) mice. ****p<0.0001, 1-way ANOVA. (j) Quantification in the DG of VCAM1+Lectin+ stained brain vasculature of young isochronic or heterochronic parabionts 5 weeks after surgery. Representative Images shown in Supplemental Figure 1f. **p<0.0006, 1-way ANOVA. n= 8-13 mice/group from two independent experiments.

### Circulating and membrane bound VCAM1 increase with age and in young mice exposed to aged blood

To identify proteins changing with human aging and possibly associated with the BBB and cerebrovascular function, we searched for those involved in vascular function in the healthy aging control group in a previously published plasma proteomic dataset from our lab ^10^. Concentrations of 31 factors correlated significantly with age (**Fig. 1d**, **Supplementary Table 2**, p<0.05). Of these, 8 are expressed in mouse BECs at the transcriptional level **(Supplementary Table 2**; **Supplementary Fig. 1d-e**), and 5 have vascular, endothelial, or angiogenic related functions **(Supplementary Table 2**; GeneCards). Among the proteins expressed in or related to the vasculature, sVCAM1 correlated most strongly with age (**Fig. 1e**). We confirmed this age-dependent increase in plasma sVCAM1 in an independent cohort of healthy individuals with ELISA (**Fig. 1f**). VCAM1, a member of the immunoglobulin superfamily, is upregulated on endothelium in response to inflammation throughout the body where it facilitates leukocyte tethering through the integrin receptor a4p1 (also known as VLA-4) and transmigration into tissues ^24,25^. VCAM1 is shed constitutively through proteolytic cleavage by the membrane-bound metalloproteinase ADAM17, resulting in high quantities of plasma sVCAM1 ^26,27^. Similar to humans, mice show a significant increase in plasma sVCAM1 with more advanced age (19-month-old) that is not seen in middle age (10–12-month-old; **Fig. 1g**).

This increase in sVCAM1 in plasma is associated with a significant increase in VCAM1 expression in lectin and Meca99 immunoreactive cells, markers of cerebral blood vessels, in the aged mouse dentate gyrus (**Fig.1 h-i**) ^28,29^. This increase was confirmed by quantifying CD31+VCAM1+ BECs in the cortex and hippocampi of young and aged mice (**Supplemental Fig. 1f-j**). Interestingly, exposure to aged blood through heterochronic parabiosis induced a similar increase in VCAM1 immunoreactivity in young mice (**Fig. 1j**; **Supplemental Fig. 1l**) and a concomitant increase in sVCAM1 in plasma **(Supplementary Fig. 1k**). In bulk population of BECs, mRNA expression of molecules involved in leukocyte adhesion were low or undetectable **(Supplementary Fig. 1d**). At the protein level, VCAM1 expression is visible in a small percentage of BECs (**Supplemental Fig. 1f-j**), suggesting that different populations and regions of BECs at the BBB respond uniquely to an aged systemic milieu.

Considering the heterogeneity of the BBB and the low percentage of BECs that express VCAM1 (**Fig. 1h-j**; **Supplemental Fig. 1f-j**; **Fig. 2a**), we performed single cell RNAseq on VCAM1-enriched BECs to characterize the unique molecular and phenotypic nature of rare VCAM1+ BECs. Full-length single-cell RNA-seq was performed on 160 and 112 BECs isolated from the hippocampi of pooled young and aged mice, respectively. Half of isolated BECs were VCAM1^high^ based on retroorbital labeling of VCAM1+BECs with a fluorescently labeled anti-VCAM1 mAb prior to mice perfusion and gating for collection of VCAM1^high^ single cells in 50% of wells to enrich for this rare population **(Supplementary Fig. 1f-j**; Methods). All isolated cells expressed both pan-endothelial *(Pecam1, Cldn5)* and BBB-specific markers *(Slcolcl, Slc2a1, Abcb1a)* (**Fig. 2b**; ^30–32^). Furthermore, we verified that VCAM1 protein levels correlate with *Vcam1* mRNA (**Fig. 2c**; **Supplementary Fig. 2a**). Unsupervised clustering in principal component space using the top 2500 correlated and anti-correlated genes revealed three molecularly distinct populations (**Fig. 2d**). Interestingly, none of the clusters were significantly enriched for young or aged cells, indicating that strong sources of variation exist besides age that are resulting in transcriptional heterogeneity between the BEC subpopulations (**Fig. 2e**). In spite of this, a direct comparison of *Vcam1* expression levels within the isolated VCAM1^high^ CD31+ BECs showed significantly higher *Vcam1* mRNA levels in aged BECs compared to young (**Fig. 2f**).

**Figure 2.**
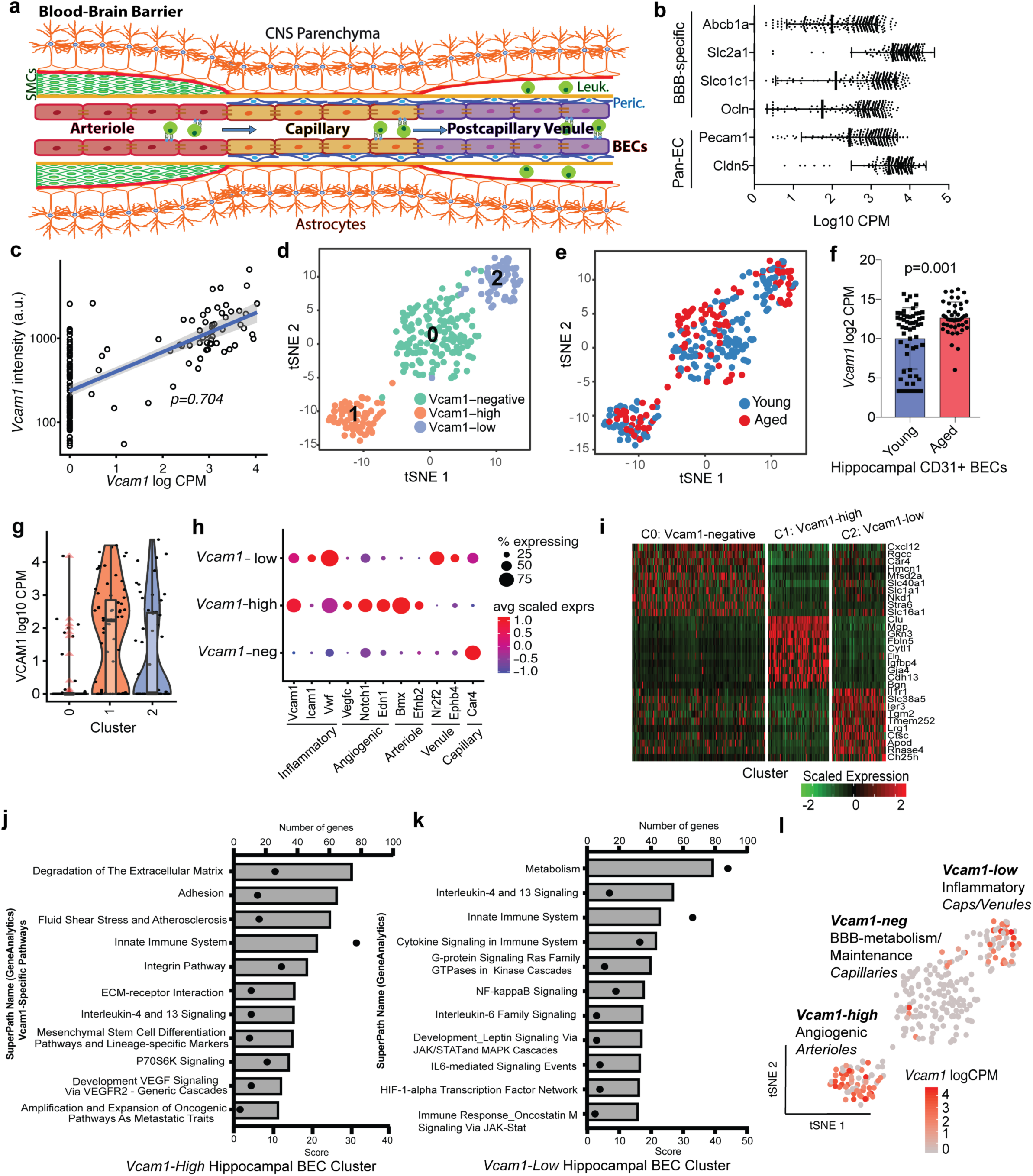
Single cell RNASeq of VCAM1 Enriched young and aged BECs reveals two unique subclusters enriched in inflammatory and cell cycle inhibitor pathways. (a) Schematic of the Blood-brain barrier (BBB). Nutrient-rich, oxygenated blood is pumped into the brain through cerebral arterioles which are protected and supported by smooth muscle cells (SMCs) that cover the endothelium and form a basement membrane layered by astrocytic end-feet of the brain parenchyma. The blood is transferred to highly specialized capillaries, which are comprised of brain endothelial cells that form unique tight junctions and are wrapped by pericytes (Peric.) within the endothelial basement membrane, which is then covered by astrocytic end-feet. BBB capillaries are the site of controlled transport of fluids and solutes into the CNS. Immuno-surveillance and occasional extravasation of leukocytes (Leuk.) into the CNS parenchyma occurs at the level of postcapillary venules, the vascular segments into which blood flows after passing through the capillaries. Postcapillary Venules contain enlarged perivascular space between the endothelial and astrocytic basement membranes where occasional immune cells can reside.^62,63^ (b) Boxplot of expression levels of classical pan-endothelial and BBB-specific transcripts (c) Validation of the correlation (Spearman’s rho = 0.704) between protein and mRNA levels of 77 single BECs sorted from both *Vcam1+ and Vcam1-*gates. Scatterplot of *Vcam1* fluorescence intensity as measured by FACs and corresponding transcript counts (per million). (d) Unbiased clustering of 112 aged and 160 young hippocampal BECs using whole transcriptome and visualization with tSNE reveals 3 molecularly distinct BEC populations. (e) tSNE visualization colored by cell identity (aged vs. young) (f) Comparison of *Vcam1* expression levels in young and aged hippocampal CD31+ BECs collected from the VCAM1+ gate during FACs sorting (error bars = SD) (g) Violin plots of *Vcam1* reveal differing levels of the transcript in each of the cell clusters. (h) Dotplot comparing the expression (scaled transcript counts and percent of population expressing) of various classical inflammatory, pro-angiogenic, arteriolar, venular and capillary markers between the three clusters (Cluster 0: Vcam1-negative, Cluster 1: Vcam1-high, Cluster 2: *Vcam1*-low). (i) Heatmap of the scaled expression of the top 10 enriched genes (differentially expressed with p>0.05, Mann-Whitney test) in each cluster. Genes are ranked by highest log-fold change when compared to all other cells. (j) GeneAnalytics (GSEA Package)-Brain Endothelial Cell Pathway analysis of the *Vcam1-high* cluster. The top 11 GO pathways containing *Vcam1* are highlighted here, along with the number of genes in each pathway enriched in this BEC cluster and the score assigned to the pathways. (k) GeneAnalytics (GSEA Package)-Brain Endothelial Cell Pathway analysis of the *Vcam1-low* cluster. The top 11 GO pathways are highlighted here, along with the number of genes in each pathway enriched in this BEC cluster and the score assigned to the pathways. (l) tSNE visualization colored by *Vcam1* expression levels. Clusters are further annotated by their putative functional-phenotype and vessel segmental identity.

Among the three unique clusters, we identified one largely *Vcam1* negative population characterized by genes relating to BBB-maintenance, metabolism and the capillary phenotype (**Fig. 2g-l**). Interestingly, the remaining two clusters were both Vcam1+, but molecularly distinct, with one expressing slightly higher *Vcam1* levels (C1) than the other (C2) (**Fig. 2g**). Using a biased classification method with known markers of the 3 main vessel types found in the BBB (**Fig. 2h**; ^30–32^), we found the Vcam1-low cluster to express significantly higher levels of pro-inflammatory genes *(Icam1, Vwf* among others) and post-capillary venule markers *(Nr2f2, Ephb4)*, while the Vcam1-high cluster expressed pro-angiogenic markers *(Vegfc, Notch1, Edn1*, among others) and the arteriolar classification *(Bmx, Efnb2)* (**Fig. 2h**, **Supplementary Fig. 2b-e**; GeneCards; ^30–32^).

We further applied the Mann-Whitney test to find differentially expressed markers between the three BEC subpopulations in an unbiased manner (**Fig. 2i**, **Supplementary Fig.2f**). Indeed, the Vcam1-low cluster was enriched with inflammatory and cytokine-signaling genes and pathways *(Il1r1, Lrg1, Ch25h*, among others), while the Vcam1-high cluster was differentially characterized by genes involved in matrix remodeling and migration *(Fbln5, Mgp)*, proliferation *(Bgn)* and angiogenesis *(Cdh13*, among others) (**Fig. 2j-l**; **Supplementary Fig.2f**; GeneCards; ^30–32^).

Given that increased expression of soluble and/or membrane VCAM1 is considered a marker of cerebrovascular inflammation and observed in encephalitis, stroke, systemic infections, and neurodegeneration in humans and mice, ^24,33^; **Supplementary Figs. 3a-f** and **7a-d**), our findings suggest that factors in aged blood are sufficient to increase cerebrovascular VCAM1 expression and thus contribute to vascular inflammation.

**Figure 3.**
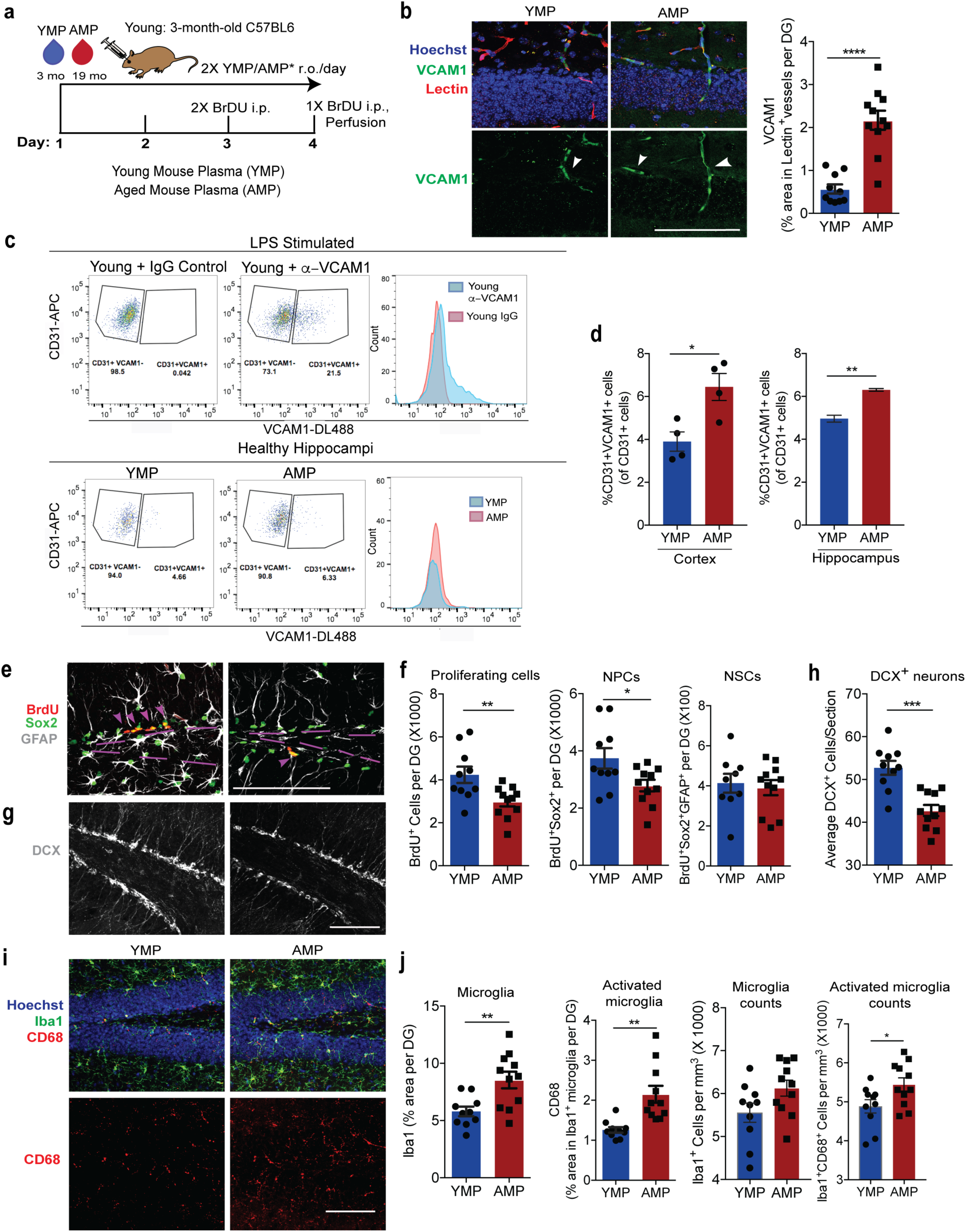
Aged blood administration into young mice activates brain vasculature and microglia and reduces hippocampal neurogenesis. (a) Schematic of experimental design. n=10-11 mice/group. (b) Representative confocal images (left) and quantification (right) of VCAM1+lectin+ in the DG. Hoechst labels cell nuclei. Arrows indicate VCAM1+ vessels. Scale bar = 100 μm. ****p<0.0002. (c) Top: Gating and histogram plots of CD31+VCAM1+ cells isolated from LPS stimulated young (3-month old) wildtype mice injected with fluorescently tagged anti-VCAM1 mAb or IgG isotype control (r.o.) 2 hours before sacrifice. Bottom: Flow gating and histogram plots of pooled (n=4 mice/plasma treatment), young or aged hippocampi isolated from plasma-injected young mice administered anti-VCAM1 mAb prior to sacrifice. Quantification (d) of CD31+VCAM1+cells isolated from healthy cortex (n=4 single mice/plasma treatment) and of 4 technical replicates of pooled hippocampi. **p<0.004., *p<0.02. (e) Representative confocal images and quantification (f) in the DG of BrdU+, Sox2+, and GFAP. Scale bar = 100 μm. Purple lines outline the SGZ and arrows indicate proliferating neural precursor cells. **p<0.01, *p<0.03. (g) Representative confocal images and quantification (h) in the DG of DCX (white). Scale bar = 100 μm. ***p<0.0002. (i) Representative confocal images and quantification (j) in the DG of CD68, Iba1, and Hoechst. Scale bar = 100 μm. **p<0.005, *p<0.04 Student’s *t-test*. All error bars indicate SEM.

### Aged plasma increases VCAM1 expression, reduces neurogenesis, and increases microglial reactivity

To determine whether soluble factors in blood, rather than other consequences of parabiosis, cause the increase in VCAM1 we added aged plasma to cultured BECs or infused it into young mice. Acutely isolated primary mouse BECs treated with aged plasma displayed significantly higher levels of VCAM1 compared to cells treated with young plasma **(Supplementary Fig. 4c-d**). Cultured Bend.3 cells, an immortalized BEC cell line expressing BBB-specific tight and adherens junctions **(Supplementary Fig. 4a**; ^34^), show a dose dependent increase in VCAM1 expression after treatment with TNF-α **(Supplementary Fig. 4b**), as reported by others ^35^. Likewise we found that Bend.3 cells treated with aged compared with young mouse or human plasma displayed significantly higher levels of VCAM1 **(Supplementary Fig. 4c-e**). Interestingly, other adhesion molecules, namely ICAM1, E-selectin, and P-selectin, were not significantly upregulated in the presence of aged plasma **(Supplementary Fig. 4f-g**). Retro-orbital infusion of young mice, twice daily for 4 days, with pooled plasma from aged, but not from young mice, (**Fig. 3a**) caused a significant increase in VCAM1 expression in lectin+ blood vessels and primary BECs (**Fig. 3b-d**), while ICAM1 was not changed **(Supplementary Fig. 4h**).

In line with previous findings (Villeda et al., 2011; Rebo et al., 2016;), aged plasma infusions reduced the numbers of BrdU+ proliferating cells overall, BrdU+Sox2+ neural precursor cells (**Fig. 3e-f**), and doublecortin (DCX)+ immature neurons (**Fig. 3g-h**) in the dentate gyrus of young mice. There was no change in the number of quiescent BrdU+Sox2+GFAP+ neural stem cells (**Fig. 3f**). Remarkably, acute injections of aged plasma also induced a prominent response in microglia, manifested in increased Iba1 immunoreactivity overall, expression of CD68 in Iba1+ cells, and numbers of CD68+Iba1+ microglia (**Fig. 3i-j**). The total number of microglia did not change with this short-term plasma treatment (**Fig. 3j**).

Similar to aged mouse plasma, repeated injections of aged human plasma induced a prominent increase in BEC-specific VCAM1 expression in young immunodeficient *NOD-scid* IL2Rg^null^ (NSG) mice over 4 days **(Supplementary Fig. 5a-b**) or 3 weeks **(Supplementary Fig. 5g-i**). The NSG mouse strain tolerates human xenografts and was used previously in our lab to demonstrate brain rejuvenating effects of human cord plasma ^12^. While NSG mice lack T and B lymphocytes and have defective natural killer cells, VLA-4 expressing innate immune cells of the myeloid lineage, including neutrophils and monocytes, are intact ^36^. Acute or long-term treatments with aged human plasma also inhibited neurogenesis in the dentate gyrus and resulted in increased microglial reactivity **(Supplementary Fig. 5c-f** and **5j-l**).

Together, these findings show that factors in aged plasma, injected over short or longer periods of time, likely conserved between mice and humans, and independent of the presence of T or B lymphocytes, are sufficient to increase VCAM1 expression in the young mouse brain, reduce hippocampal neurogenesis, and increase microglial reactivity.

### Genetic deletion of Vcam1 in BECs prevents inhibitory effects of aged plasma on neurogenesis and microglial activation

To test whether VCAM1 is simply a correlate of vascular inflammation or a possible mediator of the detrimental effects of aged plasma on the hippocampus, we deleted *Vcam1* in BECs using a Slco1c1-Cre^ERT2^ mouse – encoding tamoxifen-inducible Cre-recombinase under a brain endothelial and epithelial-specific Slco1c1 promoter ^37^. While unspecific recombination of tamoxifen-treated Slco1c1-Cre^ERT2^ mice can occur in granule neurons when crossed with a td-tomato reporter line ^37^, we did not see any expression of VCAM1 in Sox2+, GFAP+, NeuN+, or DCX+ cells in the DG of the hippocampus **(Supplementary Fig. 6a-b**). We confirmed that Vcam1^fl/fl^Slco1c1-Cre^ERT2+/-^ (Cre+) mice undergo BEC-specific *Vcam1* deletion following tamoxifen injections using a systemic LPS inflammation model **(Supplementary Fig. 7a**). Systemic LPS administration significantly upregulated BEC-specific VCAM1 in tamoxifen-treated Vcam1^fl/fl^Slco1c1-Cre^ERT2-/-^ (Cre-) control mice, but not in Cre+ littermates **(Supplementary Fig. 7a-d**). Interestingly, young mice lacking brain endothelial and epithelial *Vcam1* for 3 weeks of adulthood showed a lower baseline number of BrdU+Sox2+ neural precursor cells in the hippocampus while microglial number and activation were not affected **(Supplementary Fig. 7e-m**).

We next investigated the effects of aged plasma administration to young mice in the absence of brain endothelial and epithelial-specific *Vcam1* (**Fig. 4a**). While VCAM1 expression was absent in Cre+ brains, sVCAM1 levels remained high in the plasma of all tamoxifen-treated mice indicating unperturbed peripheral expression (**Fig. 4b-d**). Importantly, *Vcam1* deletion in Cre+ brain endothelium abrogated the detrimental effects of aged mouse plasma on hippocampal neurogenesis and microglial reactivity (**Fig. 4e-k**). Thus, Cre+ mice treated with young or aged plasma had equal levels of neurogenesis as shown by equal numbers of BrdU+, BrdU+Sox2+, and DCX+ neural precursor cell populations (**Fig. 4e-i**). Moreover, aged plasma also failed to induce microglial reactivity, indicated by the levels of CD68 in Iba1+ cells (**Fig. 4j-k**). Similar to wildtype mice (**Fig. 3**), Cre-negative control mice showed reduced neurogenesis and increased microglial activation when exposed to aged plasma (**Fig. 4e-k**). Brain endothelial and epithelial-specific Vcam1 deletion similarly prevented microglial activation and the inhibition of neurogenesis in the dentate gyri of young Cre+ mice exposed to an aged systemic milieu through long-term aged plasma administration **(Supplementary Fig. 8a-h**) or parabiosis **(Supplementary Fig. 9a-g**) with wildtype mice.

**Figure 4.**
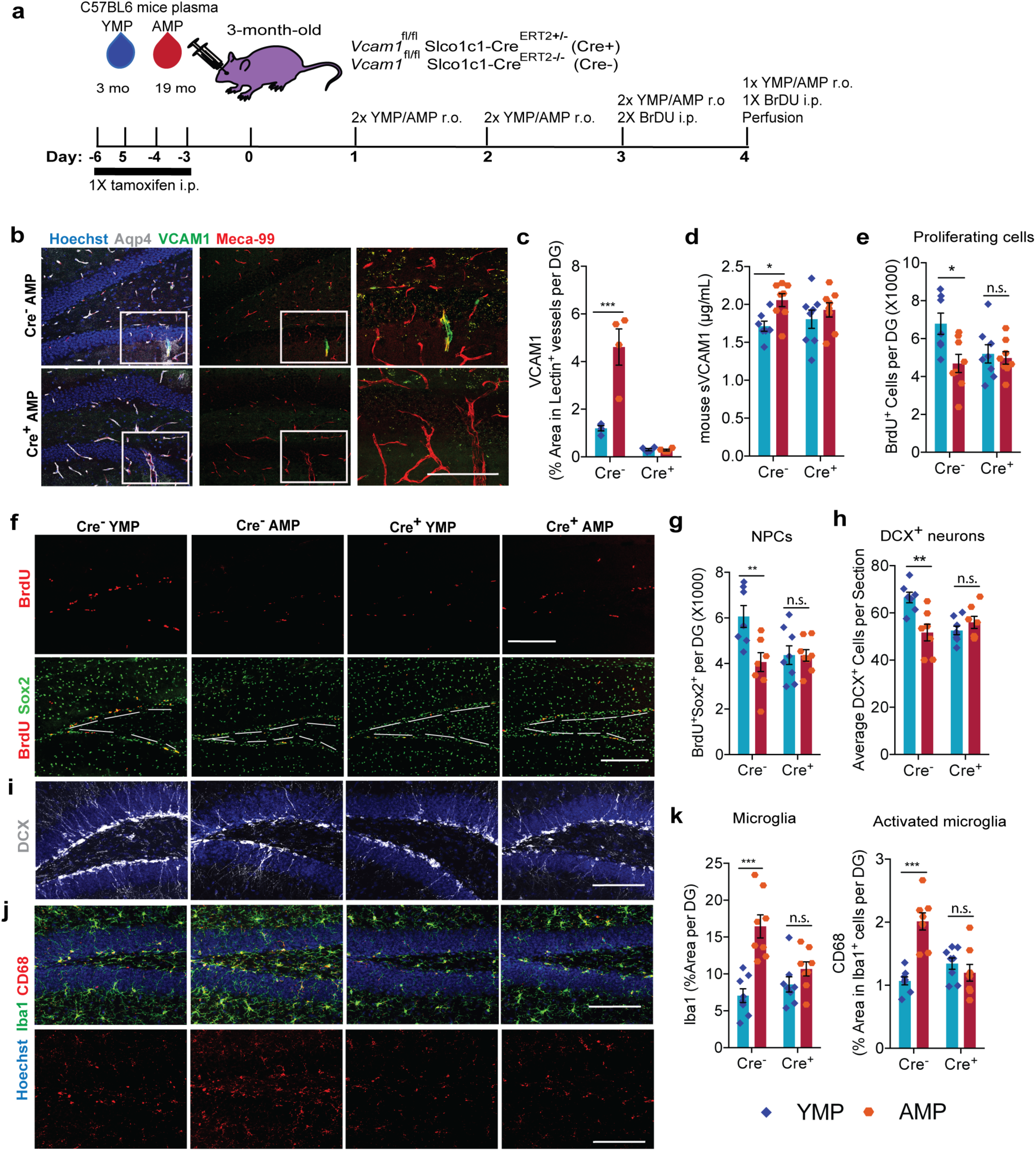
Brain endothelial and epithelial-specific *Vcam1* deletion in young mice mitigates the effects of aged plasma administration. (a) Experimental design. n=7-8 mice/group. (b) Representative confocal images in the DG of VCAM1, MECA-99, and Aqp4. Hoechst labels cell nuclei. Scale bar = 200 pm for merged images and scale bar= 100 pm for the zoomed VCAM1 and MECA-99 merged images outlined with white squares. (c) Quantification of VCAM1+ lectin+ vasculature ***p<0.004. (d) Mouse sVCAM1 ELISA of plasma samples. *p<0.03. (e) BrdU quantification and representative confocal images (f) and BrdU+Sox2+ quantification (g) in the DG of brain sections immunostained for BrdU and Sox2. White dotted lines outline the SGZ; Scale bar = 200 μm. *p<0.03, **p<0.02. (h) DCX+ quantification and representative confocal images (i) in the DG. Hoechst labels cell nuclei. Scale bar = 100 μm. **p<0.002. (j) Representative confocal images and quantification (k) from the DG of CD68 and Iba1. Hoechst labels cell nuclei. Scale bar = 100 μm. ***p<0.0009, **p<0.007, 2-way ANOVA with Tukey’s *post-hoc* test. All error bars represent SEM.

Importantly, depletion of sVCAM1 from aged human plasma prior to in vivo administration (**Fig. 5a-d**) did not significantly change its adverse effects on neurogenesis and microglial reactivity in young mice. Specifically, both IgG and anti-sVCAM1 depleted aged human plasma upregulated mouse VCAM1 on brain endothelium (**Fig. 5e-f**), reduced hippocampal BrdU+Sox2+ neural precursor cells and DCX+ immature neurons (**Fig. 5g-i**) and increased microglial activation (**Fig. 5j**). Thus, high levels of circulating, soluble VCAM1 do not drive aging phenotypes in the young brain. Collectively, these findings show that expression of VCAM1 in BECs is both necessary for baseline maintenance of neurogenesis and for mediating detrimental effects of an aged systemic environment.

**Figure 5.**
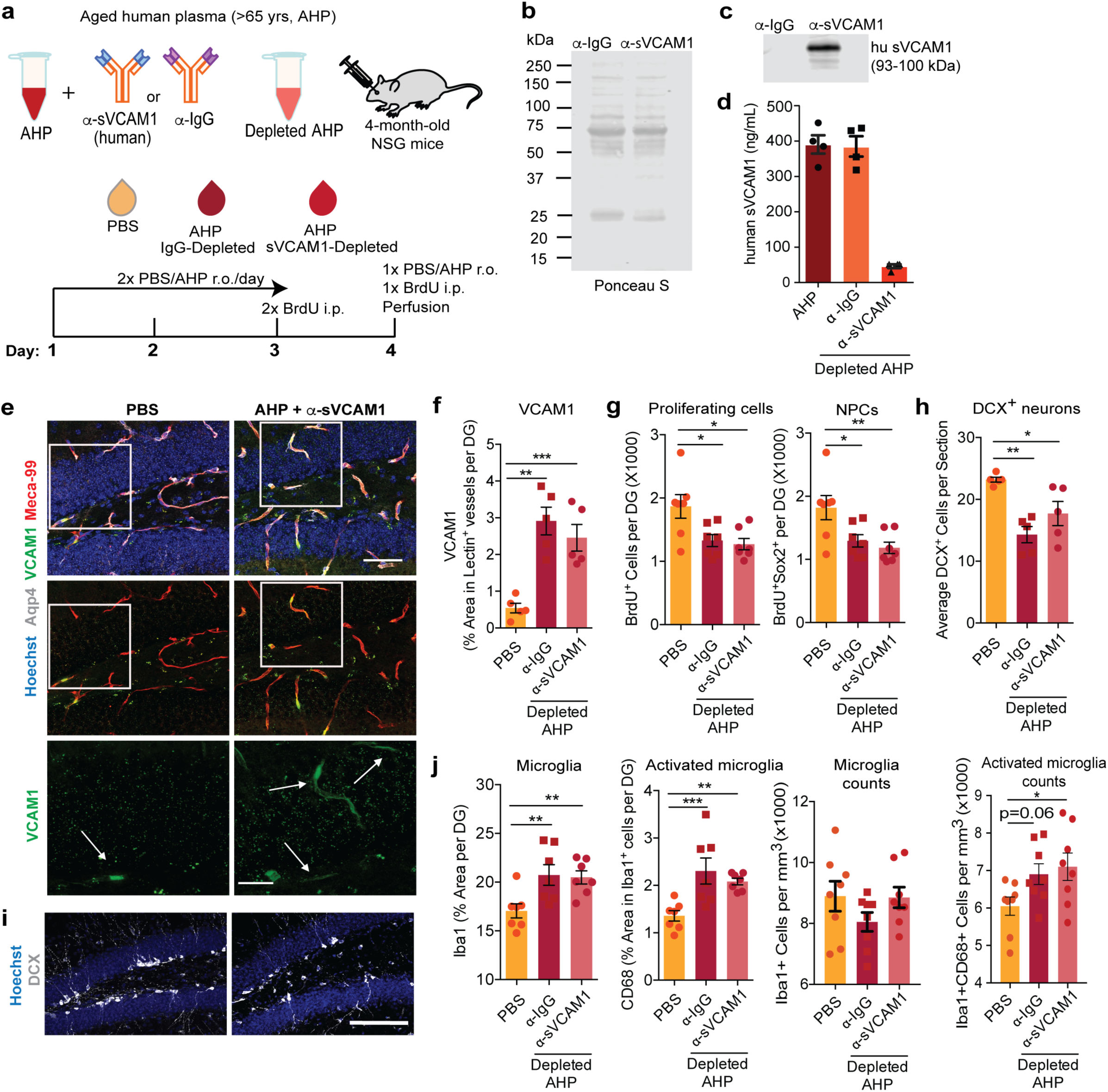
High levels of circulating sVCAM1 does not contribute to inhibitory and pro-inflammatory effects of aged plasma administration in young animals. (a) Experimental design. n=7-8 mice/group. (b) Ponceau S stain showing total protein pull-down from plasma by both IgG and anti-VCAM1 mAb conjugated beads. (c) Western blot showing human sVCAMI (93 kDa) pulled down during immunodepletion. (d) Human sVCAMI ELISA of depleted plasma. (e) Representative confocal images and quantification (f) in the DG of VCAM1, MECA-99, and Aqp4. Hoechst labels cell nuclei. Scale bar = 50 pm for merged images and scale bar= 20 pm for the 4x zoomed single channel VCAM1 images outlined with white squares. Arrows indicate VCAM1+ vessels. n=5 mice/group. ***p<0.001. (g) Quantification of the total number of BrdU+ and BrdU+Sox2+ co-labeled neural progenitor cells in the DG of immunostained sections. *p<0.009. (h) Quantification and representative confocal images (i) of the DG for DCX and Hoechst to label cell nuclei. Scale bar = 100 μm. n= 5 mice/group **p<0.003. (j) Quantification of the Iba1+ and CD68+ staining from confocal images in the DG. *p<0.05, **p<0.01, ***p<0.004. 1-way ANOVA with Tukey’s multiple comparisons *post-hoc* test. All error bars represent SEM.

### Monoclonal VCAM1 antibody prevents detrimental effects of aged plasma and reverses age-related impairments

In the CNS, VCAM1 was shown to be upregulated during epileptic seizures, and blockade with an antibody significantly reduced seizures in a mouse model ^20^. Leukocytes bind to VCAM1 primarily through a4R1 integrin, also known as VLA-4 ^38^. A monoclonal antibody that targets VLA-4 is approved for use in the clinic to treat multiple sclerosis (MS) and Crohn’s Disease ^39^. To determine whether blocking VCAM1-VLA4 interaction would mimic the effects of genetic VCAM1 deletion, we administered an anti-VCAM1 or an IgG isotype control antibody systemically into young mice treated with aged plasma (**Fig. 6a**). This well-characterized anti-VCAM1 antibody binds to immunoglobulin domains 1 and 4 of the extracellular domain of the protein and prevents leukocyte tethering^40^. We find systemic administration of fluorescently conjugated anti-VCAM1 antibody decorates cerebral blood vessels in aged mice (**Fig. 1h**) and in young NSG mice injected with aged human plasma (**Fig. 5e**). While anti-VCAM1 treatment did not affect the increase in VCAM1 expression in brain endothelium following aged plasma infusion (**Fig. 6b,e**), it completely prevented the inhibitory effects of aged plasma on neurogenesis (**Fig. 6c,f**). Anti-VCAM1 antibody treatment also prevented the increase in reactive microglia following aged plasma infusion (**Fig. 6d,g**). In contrast, in PBS-treated control mice, anti-VCAM1 treatment had no effects on VCAM1 expression, neurogenesis, or microglial reactivity parameters measured. Anti-VCAM1 antibody treatment similarly prevented the inhibitory effects of aged human plasma administration in young NSG mice treated over the course of 3 weeks **(Supplementary Fig. 10a-g**).

**Figure 6.**
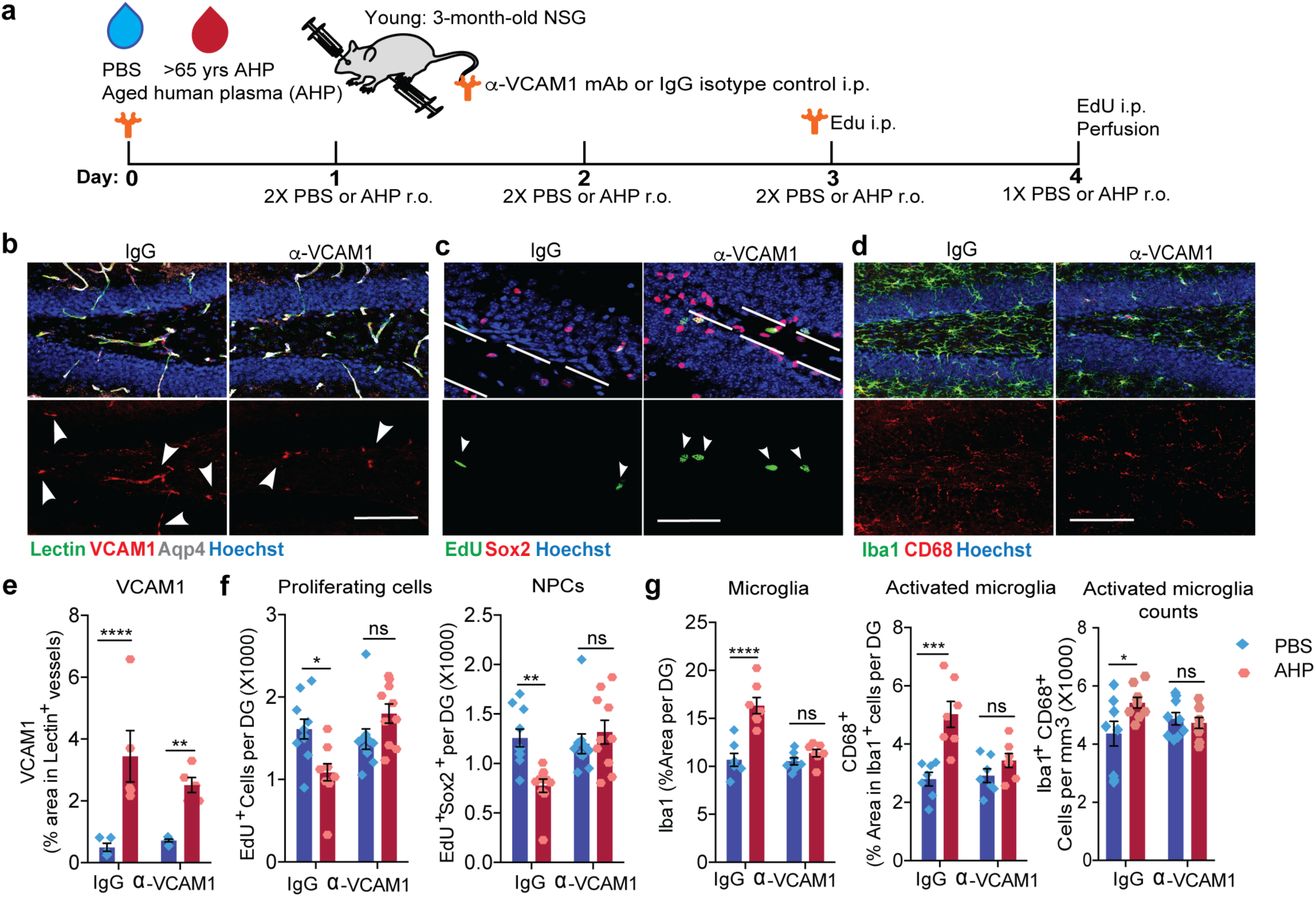
Anti-VCAM1 antibody prevents inhibitory effects of aged plasma administration in young mice. (a) Experimental design. n=10 mice/group. (b) Representative confocal images and quantification (e) (n= 5 mice/group) in the DG of VCAM1, lectin, and Aqp4. Hoechst labels cell nuclei. White arrows point to VCAM1+ vessels. Scale bar = 100 μm. ****p<0.0001, 2-way ANOVA. (c) Representative confocal images and quantification (f) in the DG of EdU and Sox2. Hoechst labels cell nuclei. Arrows indicate proliferating neural precursor cells. The SGZ is outlined with white lines. Scale bar = 50 μm. *p<0.02, 2-way ANOVA. (d) Representative confocal images and quantification (g) in the DG of CD68 and Iba1. Hoechst labels cell nuclei. Scale bar = 100 μm. ****p<0.0007, ***p<0.002, *p<0.04, 2-way ANOVA.

The above results indicate that the VCAM1 increase in brain endothelium induced by aged plasma may mediate, in part, its detrimental effects on hippocampal neurogenesis and microglial reactivity. To determine if VCAM1 induced as a result of normal aging may mediate similar negative effects on the brain, we treated aged wildtype mice with anti-VCAM1 antibody (**Fig. 7a**). Strikingly, we observed a general increase in BrdU+ proliferating cells as well as in BrdU+Sox2+ neural precursor cells in mice that received anti-VCAM1 antibody for 3 weeks (**Fig. 7b,d**). We also observed a significant reduction in the number of Iba1+CD68+ activated microglia (**Fig. 7c,e**). Anti-VCAM1 antibody treatment similarly promoted neural precursor cell proliferation and reduced microglial activation in aged female mice which, like male mice, show age-related increases in levels of soluble and BEC-specific VCAM1 **(Supplementary Fig. 10h-l**). To determine if the VCAM1 ligand VLA-4 is similarly involved in regulating age-related neurogenesis and microglial reactivity, we treated aged wildtype mice with anti-VLA-4 antibody, which did not affect VCAM1 levels in the hippocampus (**Fig. 7f-g,i**). While we observed a reduction in Iba1+CD68+ activated microglia as well, neural progenitor proliferation was unaffected (**Fig. 7h,j-k**). In summary, systemic antibody blockade of VCAM1 prevented the inhibitory effects of an aged systemic milieu on neurogenesis and microglial reactivity and blockade of VLA-4 mimicked these effects partially. All in vivo experiments are summarized in **Supplementary Table 3**.

**Figure 7.**
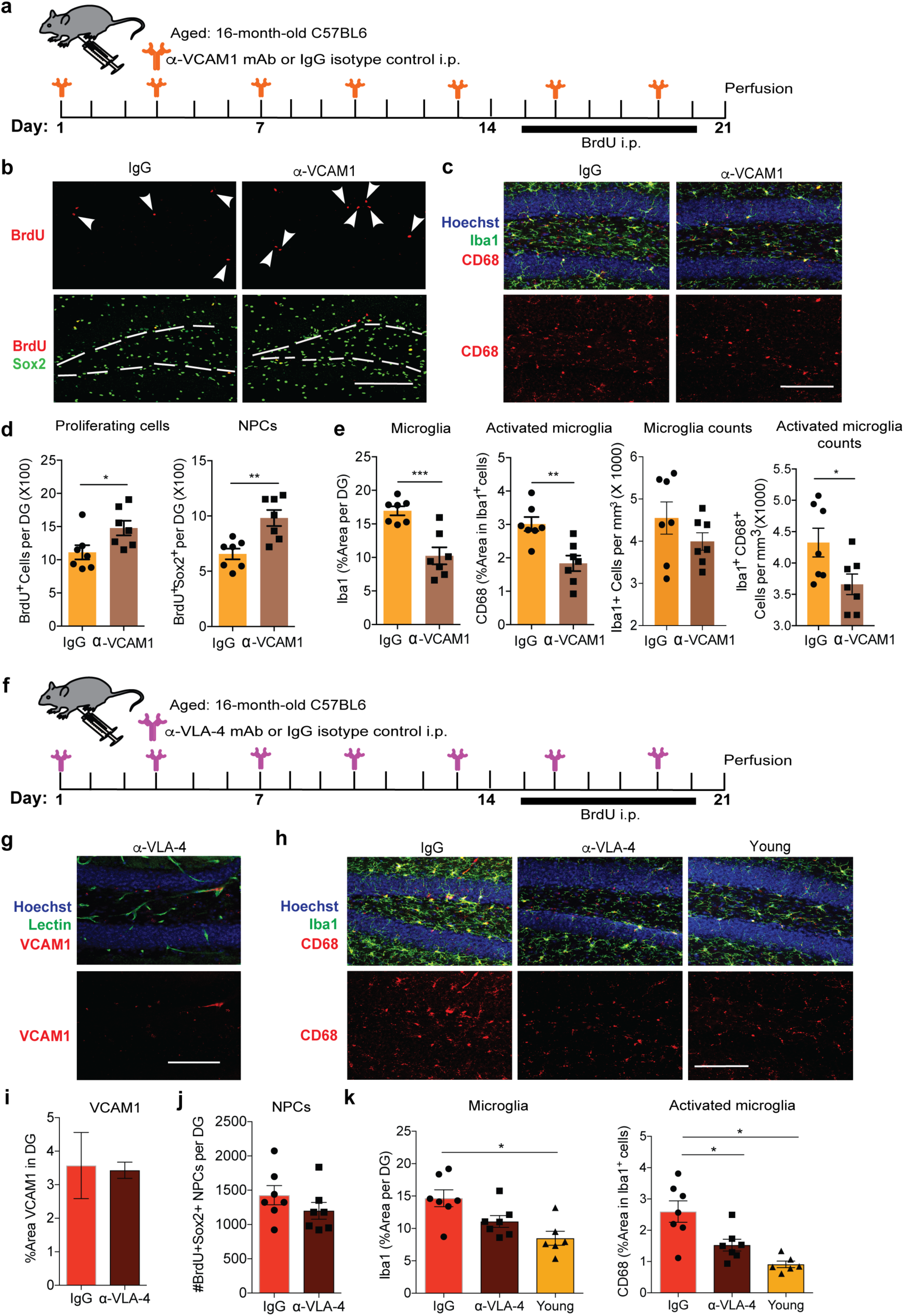
Anti-VCAM1 and anti-VLA-4 antibody rejuvenates aged brains. (a) Experimental design for anti-VCAM1. n=7 mice/group. (b) Representative confocal images and quantification (d) in the DG of BrdU and Sox2. Arrows indicate proliferating neural precursor cells. The white lines outline the SGZ. Scale bar = 100 μm. *p<0.04, **p<0.003, Student’s *t-test*. (c) Representative confocal images and quantification (e) in the DG of CD68, Iba1, and Hoechst. Scale bar = 100 μm. p***<0006, **p<0.02, *p<0.04, Student’s *t-test*. All error bars represent SEM. (f) Experimental design for anti-VLA-4. n=7 mice/group. (g) Representative confocal images and quantification (I) of VCAM1, Lectin, and Hoechst to label cell nuclei. Scale bar = 100 μm. (h) Representative confocal images and quantification (k) in the DG of CD68, Iba1, and Hoechst. Scale bar = 100 μm. *p<0.02, 1-way ANOVA. All error bars represent SEM. (j) Quantification of confocal images of the DG of NPCs co-labeled with BrdU and Sox2.

## Discussion

We describe here a mechanism of how aging and aged plasma reduce neurogenesis and increase microglial reactivity by inducing expression of VCAM1 in BECs and propose the following model: 1) factors in aged plasma induce BEC VCAM1; 2) VCAM1 facilitates tethering, but not transmigration, of leukocytes which sustain BEC inflammation; 3) inflamed VCAM1 + brain endothelium relay signals to the parenchyma to inhibit neurogenesis and activate microglia **(Supplementary Fig. 11**). We have strong support for many of these points and future work will have to test other aspects of the model and help refine or revise it.

The BBB maintains brain homeostasis by excluding most circulating macromolecules and cells from the brain parenchyma through unique tight and adherens junctions between BECs ^41^. However, the luminal side of the endothelium also provides a direct interface between peripheral organs and the brain via the circulation. Indeed, many systemic diseases result in changes in the brain vasculature and increased expression of VCAM1 has consistently been reported in atherosclerosis ^42^, colorectal cancer ^43^, metastases ^44^, lupus erythematosus ^45^, among other inflammatory diseases ^46^. On the other hand, brain diseases, including AD ^47^, MS ^48^ or epilepsy ^20^, also result in vascular changes and increased expression of VCAM1 in BECs, and elevated sVCAM1 was correlated with cognitive impairment and cerebrovascular dysfunction in 680 elderly participants ^49^.

The circulating factors mediating the observed pro-aging effects on the brain are unknown at this point, but the dialysis membrane used in our studies (cut-off 3 kDa) removes most metabolites and small molecules, including free, circulating RNAs and lipids from pooled plasma prior to injection into young mice. We thus think that proteins are responsible for communicating many of the circulatory signals to the brain (see methods for details). Indeed, circulating cytokines and chemokines with detrimental effects on the brain increase in blood with advanced age ^9^, and several cytokines including TNF-α, IL-1ß, and IL-4 induce expression of endothelial VCAM1 through NF-κB signaling ^50,51^. The putative aging factor is unlikely to be the shed form of VCAM1 because depletion of sVCAM1 from aged plasma did not affect its potential to inhibit neurogenesis and activate microglia (**Fig. 5**). Future studies will identify the factor(s) responsible for endothelial activation.

Circulating leukocytes enter tissues in a complex cascade of events that involves capture and tethering of the cells, rolling along the endothelium, activation, adhesion, and transmigration ^52^. VCAM1, in its canonical function, is involved in the tethering, rolling, and adhesion steps by binding leukocytes through VLA-4 ^25^, and because aging or aged plasma did not result in upregulation of ICAM1, E- or P-selectin, we postulate that leukocytes do not fully adhere to BECs nor transmigrate into the parenchyma. Indeed, during normal aging, leukocyte recruitment into the brain parenchyma is minimal or absent based on flow cytometry and immunohistochemical analysis ^9,53,54^. Likewise, heterochronic parabiosis studies using a GFP-expressing transgenic mouse showed no evidence of infiltrating GFP+ leukocytes in the brains of GFP-negative parabionts ^9^.

Despite their failure to infiltrate, it seems reasonable to conclude that leukocytes are involved in the adverse effects of aged plasma on the brain because systemic administration of blocking antibody against VCAM1 prevented the inhibition of neurogenesis and the activation of microglia (**Fig. 6** and **Supplemental Table 3**). We can rule out a dominant role of B or T lymphocytes in the observed effects because VCAM1 antibody also blocks the detrimental consequences of aged human plasma on the brain in immunodeficient NSG mice, which lack these adaptive immune cells ^36^. NSG mice retain VLA-4 expressing innate immune cells including neutrophils and other granulocytes as well as monocytes. Interactions of innate immune cells with brain endothelium contribute to seizures and the development of chronic epilepsy in a mouse model in which, as here, leukocyte transmigration is not observed ^20^. Leukocyte interactions with VCAM1 trigger signaling in the endothelium ^52,55^. Such signals may be communicated through the neurovascular unit to surrounding glia and neurons. Neutrophils have recently been shown to contribute to disease progression in AD model mice as well, although in this model the cells are thought to transmigrate and cause damage within the CNS^56^.

Maybe most surprisingly and of potential therapeutic relevance, administration of VCAM1 antibody in aged mice led to increased neurogenesis and reduced microglial reactivity compared with age-matched littermates treated with an isotype control antibody (**Fig. 7**), suggesting an age-related increase in VCAM1 may be sufficient to promote interactions with leukocytes and exert detrimental effects on the brain. In addition to VCAM1 blockade, antibodies against its leukocyte receptor, VLA-4, had beneficial effects on the aged mouse brain as well, significantly reducing microglial activation (**Fig. 7** and **Supplementary Table 3**). Interestingly, anti-VLA-4 antibody did not affect neurogenesis in aged mice, possibly because leukocytes could still tether to BECs through other pathways. Alternatively, progenitor proliferation in aged mice, already reduced to less than 10% compared with young mice ^57^, may no longer be susceptible to rescue by peripheral modulation of leukocyte binding to brain endothelium.

Intriguingly, we observe VCAM1 in only a small subset of BECs, even after LPS stimulation (**Fig. 1,3** and **Supplementary Figs.1, 7**). We propose that activated VCAM1+ BECs are specialized cells which produce and release pro-inflammatory cytokines into the brain to possibly modulate neurogenesis and microglial activity. In support of this, single cell RNAseq of isolated hippocampal BECs revealed 3 unique subpopulations of BECs: 1) *Vcam1-negative* capillaries expressing characteristic BBB genes related to maintenance and metabolism, 2) *Vcam1-high* arterioles expressing angiogenic markers, and 3) *Vcam1-low* capillaries and venules expressing slightly lower levels of *Vcam1* along with inflammatory gene transcripts (**Fig. 2** and **Supplementary Fig. 2**). Whether VCAM1 expressing BECs are specialized subtypes of endothelial cells or the result of stochastic adhesion and activation events is largely unknown. However, considering even young healthy mice contain these 2 unique VCAM1 expressing subpopulations of BECs, we hypothesize they represent specialized BECs that are primed for environmental sensing and immediate response to pathogenic stimuli. Moreover, the discovery of 2 molecularly distinct populations of VCAM1+ BECs reveals an unprecedented heterogeneity within an already rare cell population. In addition to the canonical roles of VCAM1 in inflammation, the presence of another *Vcam1-high* angiogenic population (roughly 50% of all VCAM1^high^ cells analyzed) elevates the complexity of VCAM1 in the context of the VCAM1 + BEC-neurogenesis signaling axis. For instance, it is possible that the neuro-regenerative effect of *Vcam1* deletion may involve multiple phenotypically distinct VCAM1+ BEC populations. To our knowledge, this the first detailed analyses of the BEC transcriptome of the neurogenic hippocampus at the single cell level.

In the adult mammalian brain, stem cell self-renewal and subsequent proliferation of neural progenitors is regulated by a specialized microenvironment ^58^, and changes in biochemical cues upon aging contribute significantly to age-associated neurodegeneration. VCAM1 is required under non-pathological conditions for Type B neural stem cell anchoring to the neurogenic niche of the subventricular zone (SVZ) ^59^ where it is highly expressed in endothelial cells of the lateral ventricles and epithelial cells of the choroid plexus ^59,60^. Kokovay and colleagues showed that VCAM1 expression increases in the lateral ventricles as a result of increased inflammatory cytokine signaling, leading to production of reactive oxygen species (ROS) and restriction of NSC proliferation and lineage progression in the SVZ ^59^. Antibody blockade of VCAM1 activated Type B neural stem cells to a proliferative state ^59^. Additionally, it was recently shown that VCAM1 expression in radial glial cells is necessary for embryonic neurogenesis and development of the SVZ neurogenic niche ^61^. While this seems to contradict our findings, we observed, indeed, that genetic deletion of *Vcam1* in young Cre+ VCAM1 floxed mice resulted in a reduction of baseline neurogenesis. Thus, VCAM1 appears to have dual roles in regulating adult neurogenesis, supporting a homeostatic role related to anchoring of stem cells in their niche as well as a role in aging-related inflammation which inhibits neurogenesis. In other words, genetic ablation of brain endothelial and epithelial *Vcam1* in young animals may reduce neurogenesis due to depletion of the quiescent neural stem cell population. Increased VCAM1 with aging and inflammation, on the other hand, may reduce neurogenesis by restricting NSC activation and lineage progression.

In summary, we show here that aging and aged plasma engage VCAM1 to inhibit adult neurogenesis and promote microglial reactivity and neuroinflammation, thus directly implicating the cerebrovasculature and the BBB in these age-related brain changes. While a tight BBB supports brain homeostasis, crossing the BBB remains the greatest obstacle for therapeutic interventions for the treatments of neurodegenerative diseases. The ability, therefore, to modulate brain function through the systemic milieu holds great potential for combating neurodegeneration. Indeed, our data raise the possibility that the decrease in neurogenesis and rise in neuroinflammation linked to systemic aging and disease can be ameliorated through non-invasive, systemic modulation of VCAM1 at the BBB.

## AUTHOR CONTRIBUTIONS

H. Y. and T.W.-C designed research. H.Y. and C.J.C isolated BECs, performed and analyzed flow cytometry. H.Y. and A.B. performed in vitro experiments. H.Y., A.B., J.Z., D.L., T.M. performed staining/microscopy analysis and cell counts. H.Y., A.B., and J.Z. performed ELISAs and western blots. H.Y. and C.J.C. performed tissue dissections. H.Y. and C.J.C. performed plasma injections. H.Y. and L.B. performed parabiosis. M.S. provided Slco1c1-Cre^ERT2^ breeding pair. B.L. analyzed human proteomic data. V.M., A.B., and M.B.C edited the manuscript. C.J.C. and E.C.B. helped with experimental design and edited the manuscript. C.J.C developed protocols for BEC isolation, cultivation and flow cytometry following LPS stimulation. H.Y. and E.B. performed bulk RNAseq experiment. H.Y., M.B.C, and D.L. performed single cell RNAseq experiment. SRQ supervised scRNAseq data collection and analysis and reviewed the manuscript. H.Y., R.V.N. and B.L. and E.B. analyzed bulk transcriptomic data. M.B.C and H.Y. analyzed single cell transcriptomic data. H.Y. analyzed data and generated the figures. H.Y. and T.W.-C wrote the manuscript. T.W.-C. supervised the study.

## ACKNOWLEDGEMENTS

We thank Lusijah Sutherland, PhD, and Corey Cain, PhD, for managing the core flow cytometry facility at the VA in Palo Alto and providing H.Y. and C.J.C. training on the instruments; Husein Hadeiba, PhD (Scientist at VA Palo Alto Health Care System) for his experimental advice, staining and analysis of PBMCs and thoughtful discussion; Oscar Leyva for assistance in staining/microscopy analysis for the experiment shown in Supplemental Figure 4. We would also like to thank Ryan Watts, PhD and Nga Bien-Ly, PhD for sharing a BEC isolation protocol used for RNAseq. This work was funded by the Department of Veterans Affairs (T.W.-C.), the National Institute on Aging (SPO: 116650; 1F32AG051330-01A1 to H.Y., R01-AG045034 and DP1-AG053015 to T.W.-C.), the NOMIS Foundation (T.W.-C.), The Glenn Foundation for Aging Research (T.W.-C), a SPARK grant to H.Y. through the Stanford Clinical and Translational Science Award (CTSA) to Spectrum (UL1 TR001085), the National Institutes of Health (R01-GM37734 and R37-AI047822 to E.C.B), the Stanford Institute for Immunity, Transplantation and Infection (C.J.C.), and the Edinger Institute (C.J.C.). The CTSA program is led by the National Center for Advancing Translational Sciences (NCATS) at the National Institutes of Health (NIH).

---

**Supplemental Table 1:**
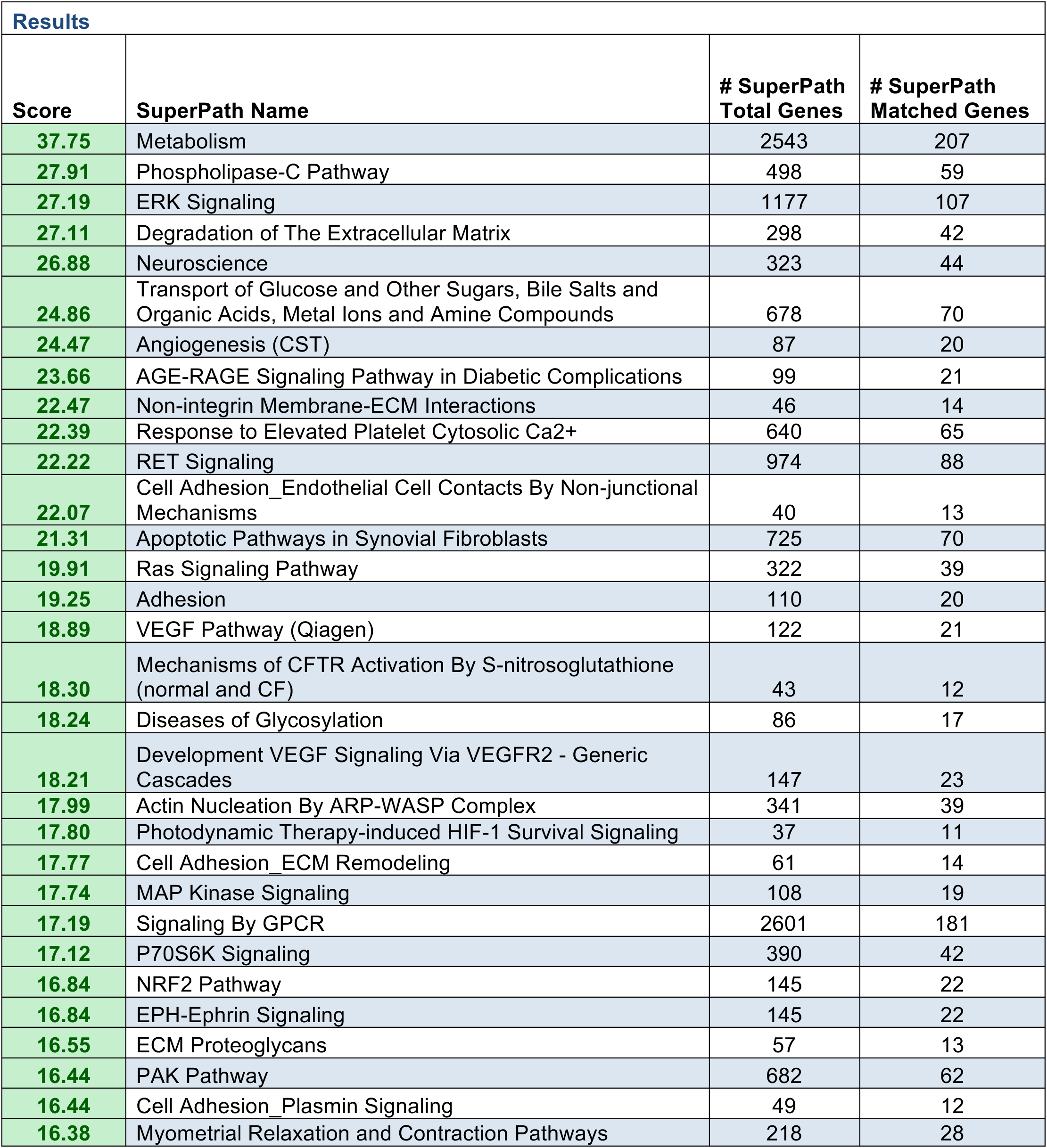
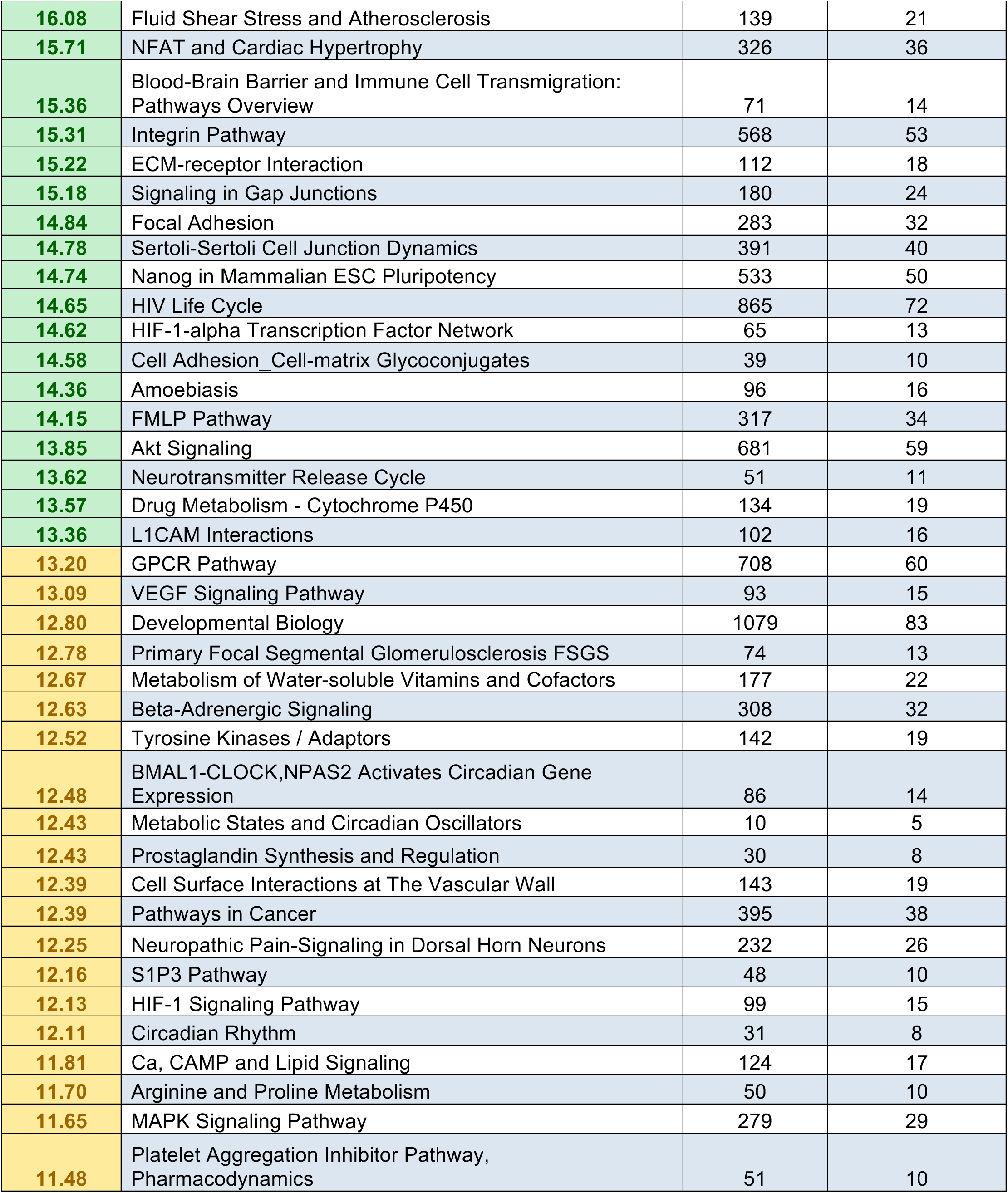
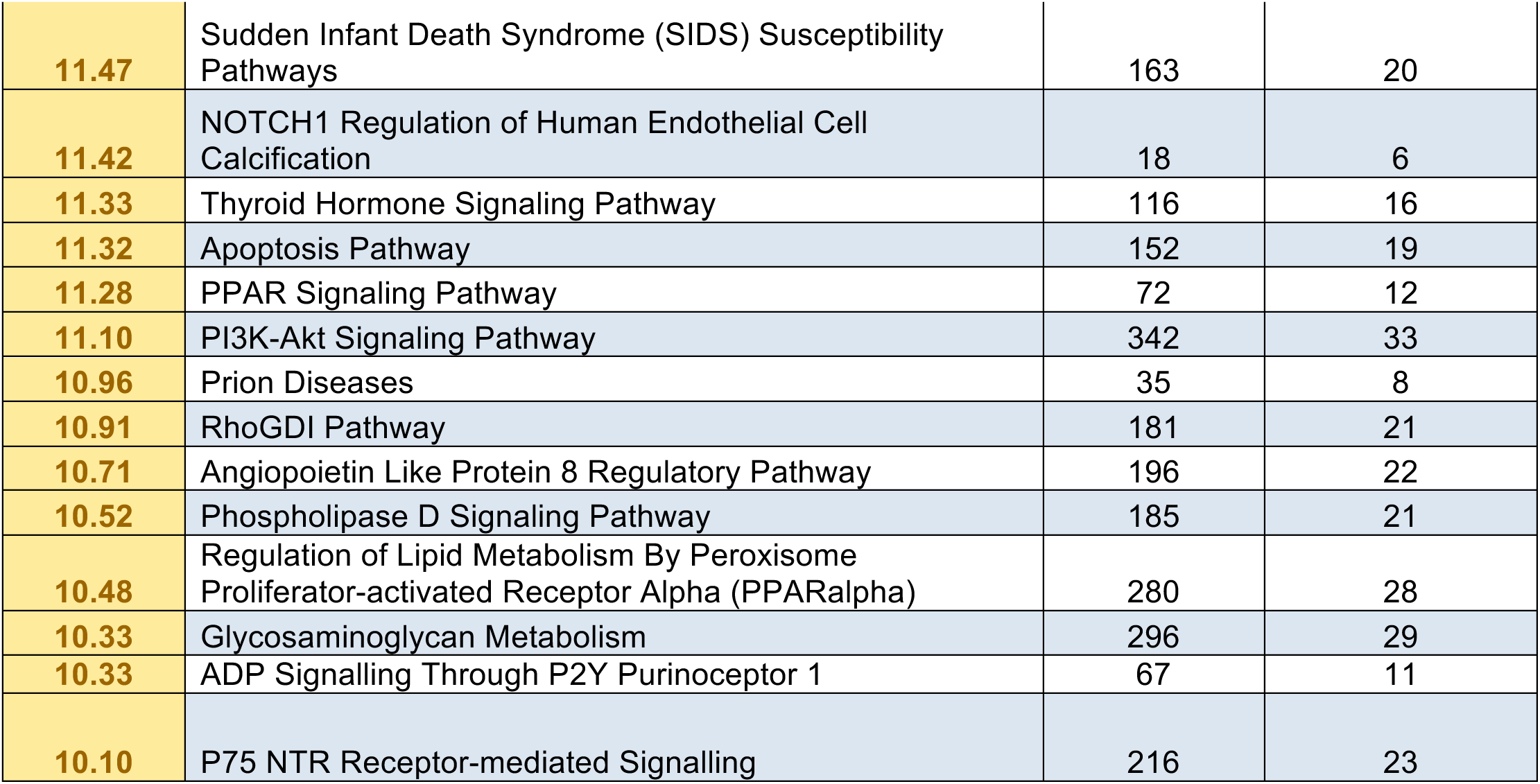
Bulk RNAseq of Young versus Aged Pooled Hippocampal and Cortical BECs: SuperPathways Analysis. Canonical pathways that are differentially regulated in aged BECs identified by GeneAnalytics (GSEA Package). *Related to Figure 1*

| **Results** |  |  |  |
| --- | --- | --- | --- |
| **Score** | **SuperPath Name** | **# SuperPath Total Genes** | **# SuperPath Matched Genes** |
| **37.75** | Metabolism | 2543 | 207 |
| **27.91** | Phospholipase-C Pathway | 498 | 59 |
| **27.19** | ERK Signaling | 1177 | 107 |
| **27.11** | Degradation of The Extracellular Matrix | 298 | 42 |
| **26.88** | Neuroscience | 323 | 44 |
| **24.86** | Transport of Glucose and Other Sugars, Bile Salts and Organic Acids, Metal Ions and Amine Compounds | 678 | 70 |
| **24.47** | Angiogenesis (CST) | 87 | 20 |
| **23.66** | AGE-RAGE Signaling Pathway in Diabetic Complications | 99 | 21 |
| **22.47** | Non-integrin Membrane-ECM Interactions | 46 | 14 |
| **22.39** | Response to Elevated Platelet Cytosolic Ca<sup>2+</sup> | 640 | 65 |
| **22.22** | RET Signaling | 974 | 88 |
| **22.07** | Cell Adhesion_Endothelial Cell Contacts By Non-junctional Mechanisms | 40 | 13 |
| **21.31** | Apoptotic Pathways in Synovial Fibroblasts | 725 | 70 |
| **19.91** | Ras Signaling Pathway | 322 | 39 |
| **19.25** | Adhesion | 110 | 20 |
| **18.89** | VEGF Pathway (Qiagen) | 122 | 21 |
| **18.30** | Mechanisms of CFTR Activation By S-nitrosoglutathione (normal and CF) | 43 | 12 |
| **18.24** | Diseases of Glycosylation | 86 | 17 |
| **18.21** | Development VEGF Signaling Via VEGFR2 - Generic Cascades | 147 | 23 |
| **17.99** | Actin Nucleation By ARP-WASP Complex | 341 | 39 |
| **17.80** | Photodynamic Therapy-induced HIF-1 Survival Signaling | 37 | 11 |
| **17.77** | Cell Adhesion_ECM Remodeling | 61 | 14 |
| **17.74** | MAP Kinase Signaling | 108 | 19 |
| **17.19** | Signaling By GPCR | 2601 | 181 |
| **17.12** | P70S6K Signaling | 390 | 42 |
| **16.84** | NRF2 Pathway | 145 | 22 |
| **16.84** | EPH-Ephrin Signaling | 145 | 22 |
| **16.55** | ECM Proteoglycans | 57 | 13 |
| **16.44** | PAK Pathway | 682 | 62 |
| **16.44** | Cell Adhesion_Plasmin Signaling | 49 | 12 |
| **16.38** | Myometrial Relaxation and Contraction Pathways | 218 | 28 |
| **16.08** | Fluid Shear Stress and Atherosclerosis | 139 | 21 |
| **15.71** | NFAT and Cardiac Hypertrophy | 326 | 36 |
| **15.36** | Blood-Brain Barrier and Immune Cell Transmigration: Pathways Overview | 71 | 14 |
| **15.31** | Integrin Pathway | 568 | 53 |
| **15.22** | ECM-receptor Interaction | 112 | 18 |
| **15.18** | Signaling in Gap Junctions | 180 | 24 |
| **14.84** | Focal Adhesion | 283 | 32 |
| **14.78** | Sertoli-Sertoli Cell Junction Dynamics | 391 | 40 |
| **14.74** | Nanog in Mammalian ESC Pluripotency | 533 | 50 |
| **14.65** | HIV Life Cycle | 865 | 72 |
| **14.62** | HIF-1-alpha Transcription Factor Network | 65 | 13 |
| **14.58** | Cell Adhesion_Cell-matrix Glycoconjugates | 39 | 10 |
| **14.36** | Amoebiasis | 96 | 16 |
| **14.15** | FMLP Pathway | 317 | 34 |
| **13.85** | Akt Signaling | 681 | 59 |
| **13.62** | Neurotransmitter Release Cycle | 51 | 11 |
| **13.57** | Drug Metabolism - Cytochrome P450 | 134 | 19 |
| **13.36** | L1CAM Interactions | 102 | 16 |
| **13.20** | GPCR Pathway | 708 | 60 |
| **13.09** | VEGF Signaling Pathway | 93 | 15 |
| **12.80** | Developmental Biology | 1079 | 83 |
| **12.78** | Primary Focal Segmental Glomerulosclerosis FSGS | 74 | 13 |
| **12.67** | Metabolism of Water-soluble Vitamins and Cofactors | 177 | 22 |
| **12.63** | Beta-Adrenergic Signaling | 308 | 32 |
| **12.52** | Tyrosine Kinases / Adaptors | 142 | 19 |
| **12.48** | BMAL1-CLOCK,NPAS2 Activates Circadian Gene Expression | 86 | 14 |
| **12.43** | Metabolic States and Circadian Oscillators | 10 | 5 |
| **12.43** | Prostaglandin Synthesis and Regulation | 30 | 8 |
| **12.39** | Cell Surface Interactions at The Vascular Wall | 143 | 19 |
| **12.39** | Pathways in Cancer | 395 | 38 |
| **12.25** | Neuropathic Pain-Signaling in Dorsal Horn Neurons | 232 | 26 |
| **12.16** | S1P3 Pathway | 48 | 10 |
| **12.13** | HIF-1 Signaling Pathway | 99 | 15 |
| **12.11** | Circadian Rhythm | 31 | 8 |
| **11.81** | Ca, CAMP and Lipid Signaling | 124 | 17 |
| **11.70** | Arginine and Proline Metabolism | 50 | 10 |
| **11.65** | MAPK Signaling Pathway | 279 | 29 |
| **11.48** | Platelet Aggregation Inhibitor Pathway, Pharmacodynamics | 51 | 10 |
| **11.47** | Sudden Infant Death Syndrome (SIDS) Susceptibility Pathways | 163 | 20 |
| **11.42** | NOTCH1 Regulation of Human Endothelial Cell Calcification | 18 | 6 |
| **11.33** | Thyroid Hormone Signaling Pathway | 116 | 16 |
| **11.32** | Apoptosis Pathway | 152 | 19 |
| **11.28** | PPAR Signaling Pathway | 72 | 12 |
| **11.10** | PI3K-Akt Signaling Pathway | 342 | 33 |
| **10.96** | Prion Diseases | 35 | 8 |
| **10.91** | RhoGDI Pathway | 181 | 21 |
| **10.71** | Angiopoietin Like Protein 8 Regulatory Pathway | 196 | 22 |
| **10.52** | Phospholipase D Signaling Pathway | 185 | 21 |
| **10.48** | Regulation of Lipid Metabolism By Peroxisome Proliferator-activated Receptor Alpha (PPARalpha) | 280 | 28 |
| **10.33** | Glycosaminoglycan Metabolism | 296 | 29 |
| **10.33** | ADP Signalling Through P2Y Purinoceptor 1 | 67 | 11 |
| **10.10** | P75 NTR Receptor-mediated Signalling | 216 | 23 |

**Supplemental Table 2:**
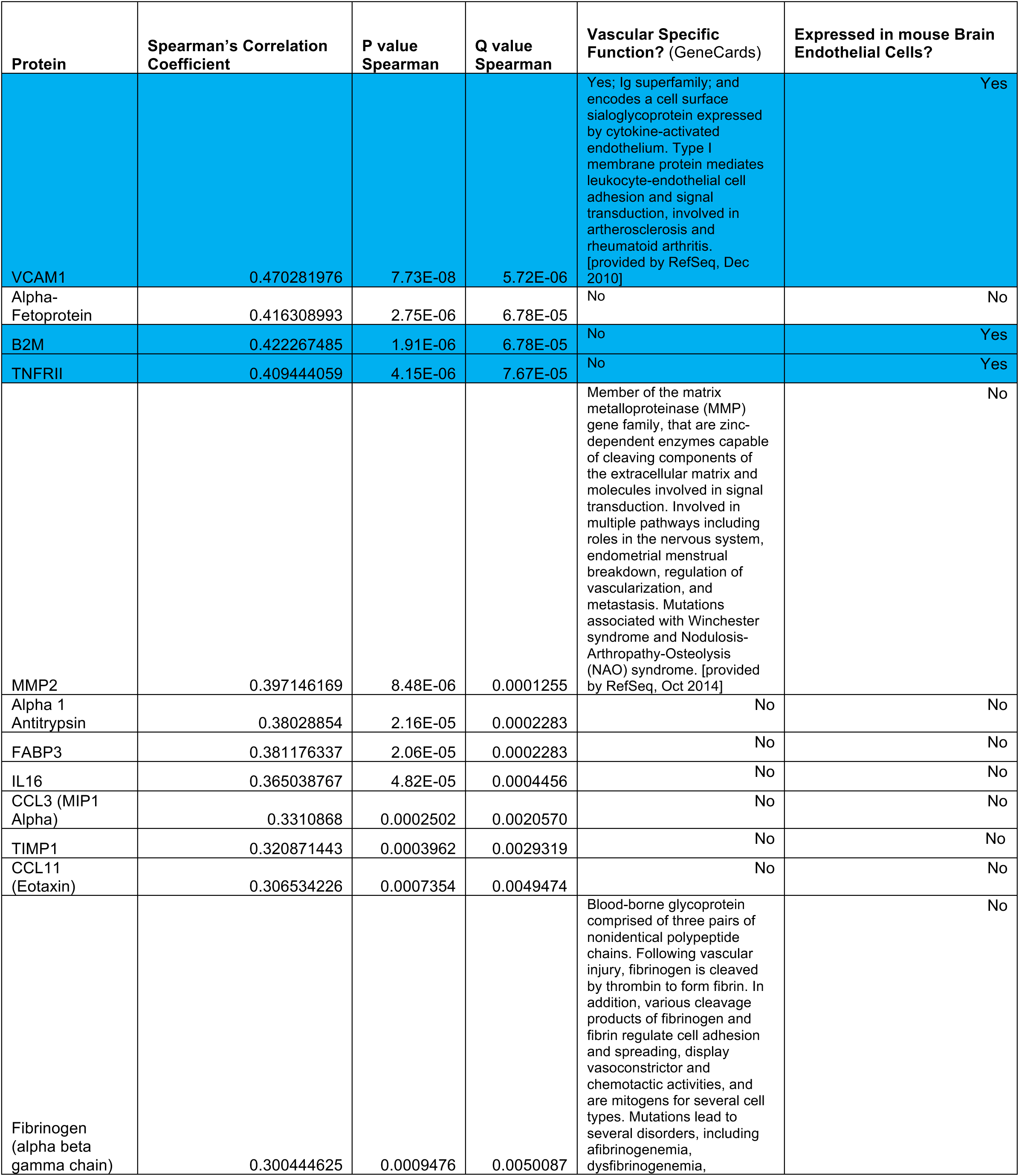
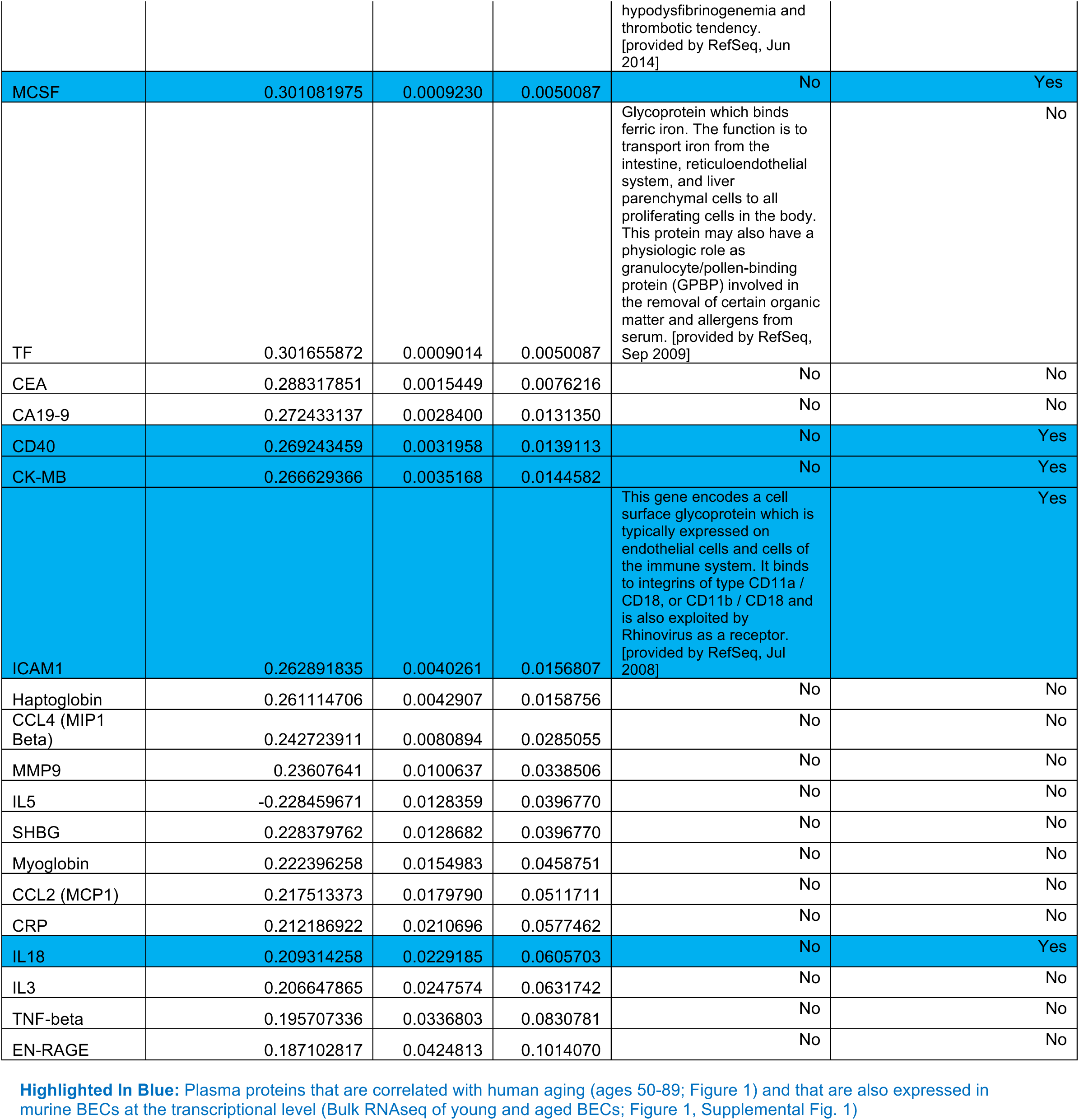
Proteomics: List of differentially regulated proteins with human aging ^10^. *Related to Figure 1*

| Protein | Spearman's Correlation Coefficient | P value Spearman | Q value Spearman | Vascular Specific Function? (GeneCards) | Expressed in mouse Brain Endothelial Cells? |
| --- | --- | --- | --- | --- | --- |
| VCAM1 | 0.470281976 | 7.73E-08 | 5.72E-06 | Yes; Ig superfamily; and encodes a cell surface sialoglycoprotein expressed by cytokine-activated endothelium. Type I membrane protein mediates leukocyte-endothelial cell adhesion and signal transduction, involved in atherosclerosis and rheumatoid arthritis. [provided by RefSeq, Dec 2010] | Yes |
| Alpha-Fetoprotein | 0.416308993 | 2.75E-06 | 6.78E-05 | No | No |
| B2M | 0.422267485 | 1.91E-06 | 6.78E-05 | No | Yes |
| TNFR2 | 0.409444059 | 4.15E-06 | 7.67E-05 | No | Yes |
| MMP2 | 0.397146169 | 8.48E-06 | 0.0001255 | Member of the matrix metalloproteinase (MMP) gene family, that are zinc-dependent enzymes capable of cleaving components of the extracellular matrix and molecules involved in signal transduction. Involved in multiple pathways including roles in the nervous system, endometrial menstrual breakdown, regulation of vascularization, and metastasis. Mutations associated with Winchester syndrome and Nodulosis-Arthropathy-Osteolysis (NAO) syndrome. [provided by RefSeq, Oct 2014] | No |
| Alpha 1 Antitrypsin | 0.38028854 | 2.16E-05 | 0.0002283 | No | No |
| FABP3 | 0.381176337 | 2.06E-05 | 0.0002283 | No | No |
| IL16 | 0.365038767 | 4.82E-05 | 0.0004456 | No | No |
| CCL3 (MIP1 Alpha) | 0.3310868 | 0.0002502 | 0.0020570 | No | No |
| TIMP1 | 0.320871443 | 0.0003962 | 0.0029319 | No | No |
| CCL11 (Eotaxin) | 0.306534226 | 0.0007354 | 0.0049474 | No | No |
| Fibrinogen (alpha beta gamma chain) | 0.300444625 | 0.0009476 | 0.0050087 | Blood-borne glycoprotein comprised of three pairs of nonidentical polypeptide chains. Following vascular injury, fibrinogen is cleaved by thrombin to form fibrin. In addition, various cleavage products of fibrinogen and fibrin regulate cell adhesion and spreading, display vasoconstrictor and chemotactic activities, and are mitogens for several cell types. Mutations lead to several disorders, including afibrinogenemia, dysfibrinogenemia, hypodysfibrinogenemia and thrombotic tendency. [provided by RefSeq, Jun 2014] | No |
| MCSF | 0.301081975 | 0.0009230 | 0.0050087 | No | Yes |
| TF | 0.301655872 | 0.0009014 | 0.0050087 | Glycoprotein which binds ferric iron. The function is to transport iron from the intestine, reticuloendothelial system, and liver parenchymal cells to all proliferating cells in the body. This protein may also have a physiologic role as granulocyte/pollen-binding protein (GPBP) involved in the removal of certain organic matter and allergens from serum. [provided by RefSeq, Sep 2009] | No |
| CEA | 0.288317851 | 0.0015449 | 0.0076216 | No | No |
| CA19-9 | 0.272433137 | 0.0028400 | 0.0131350 | No | No |
| CD40 | 0.269243459 | 0.0031958 | 0.0139113 | No | Yes |
| CK-MB | 0.266629366 | 0.0035168 | 0.0144582 | No | Yes |
| ICAM1 | 0.262891835 | 0.0040261 | 0.0156807 | This gene encodes a cell surface glycoprotein which is typically expressed on endothelial cells and cells of the immune system. It binds to integrins of type CD11a / CD18, or CD11b / CD18 and is also exploited by Rhinovirus as a receptor. [provided by RefSeq, Jul 2008] | Yes |
| Haptoglobin | 0.261114706 | 0.0042907 | 0.0158756 | No | No |
| CCL4 (MIP1 Beta) | 0.242723911 | 0.0080894 | 0.0285055 | No | No |
| MMP9 | 0.23607641 | 0.0100637 | 0.0338506 | No | No |
| IL5 | -0.228459671 | 0.0128359 | 0.0396770 | No | No |
| SHBG | 0.228379762 | 0.0128682 | 0.0396770 | No | No |
| Myoglobin | 0.222396258 | 0.0154983 | 0.0458751 | No | No |
| CCL2 (MCP1) | 0.217513373 | 0.0179790 | 0.0511711 | No | No |
| CRP | 0.212186922 | 0.0210696 | 0.0577462 | No | No |
| IL18 | 0.209314258 | 0.0229185 | 0.0605703 | No | Yes |
| IL3 | 0.206647865 | 0.0247574 | 0.0631742 | No | No |
| TNF-beta | 0.195707336 | 0.0336803 | 0.0830781 | No | No |
| EN-RAGE | 0.187102817 | 0.0424813 | 0.1014070 | No | No |

**Supplemental Table 3:**
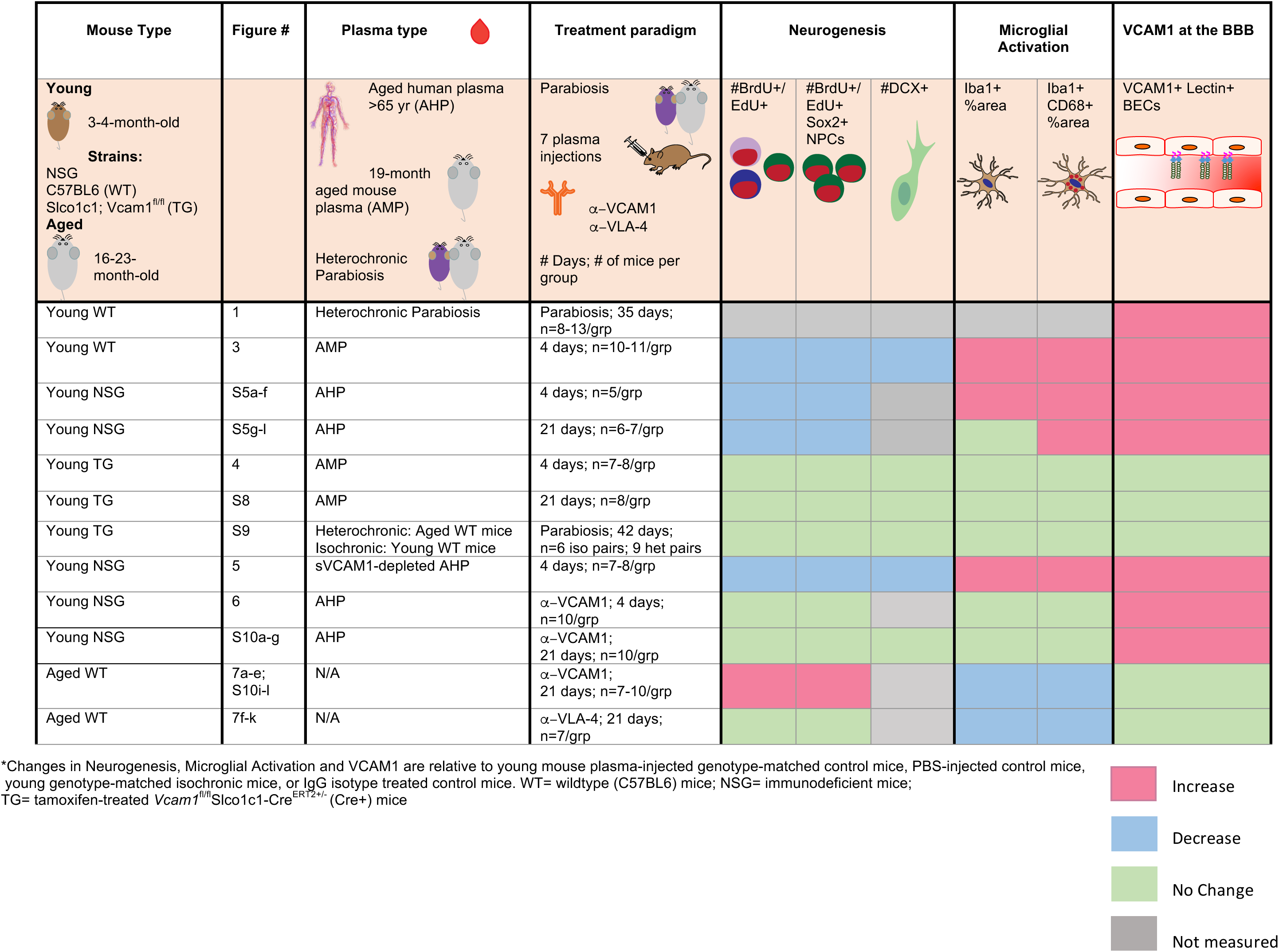
Summary of plasma treatment experiments in mice and phenotypic outcomes.

**Supplementary Figure 1.**
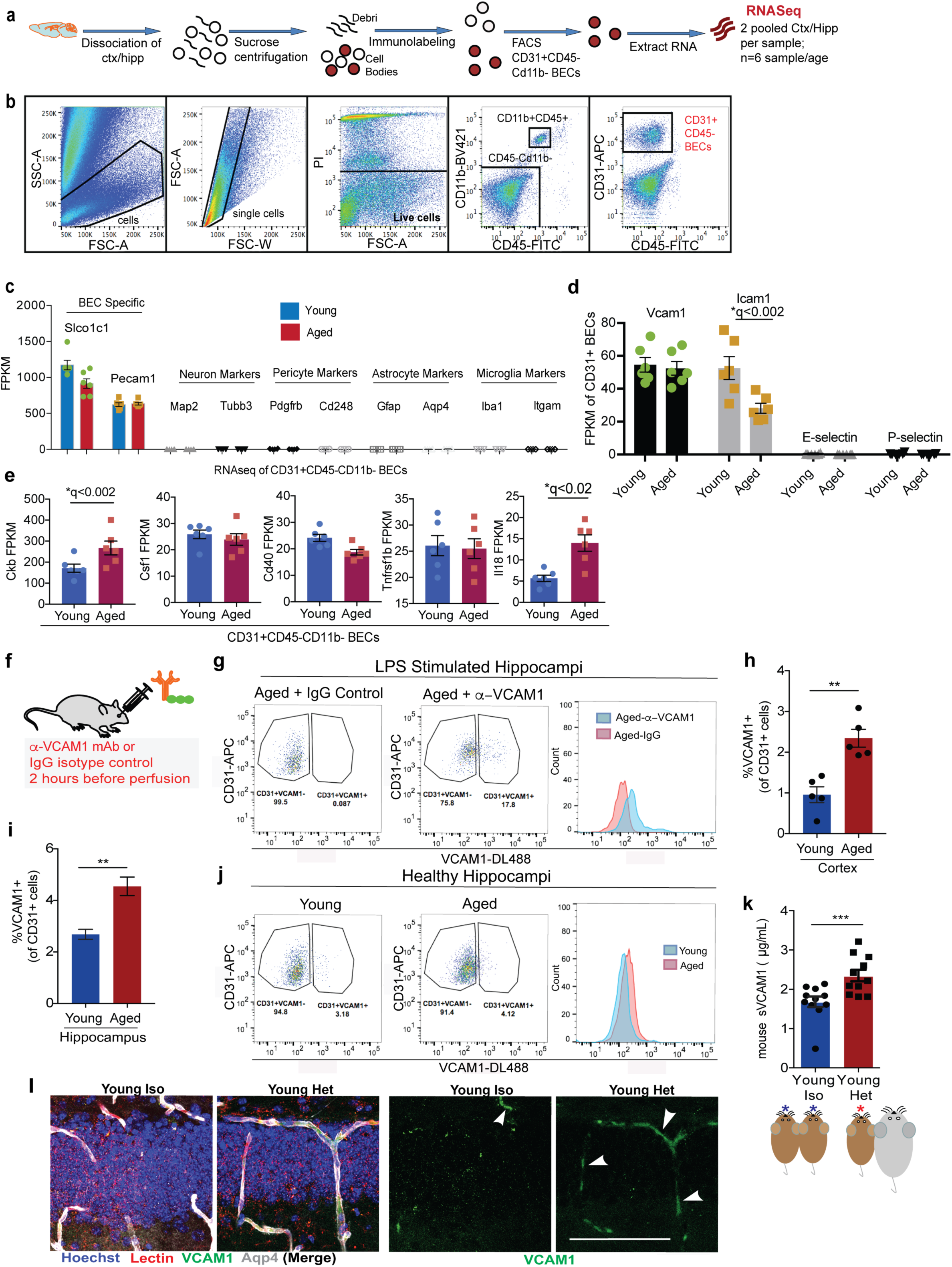
Transcriptome and proteome profiling of brain endothelial cells and plasma reveal increased inflammatory signature with aging. *Related to Figure 1* (a) Schematic of flow sorting of CD31+CD45-BECs from mouse cortex and hippocampi. Each isolated RNA sample is a pool of BECs from 2 mouse brains. (b) FACS gating strategy to isolate single BECs. PI+ dead cells were excluded. CD11b+ and CD45+ cells were gated to exclude monocytes/macrophages and microglia. CD31+Cd11b-CD45-cells were defined as the BEC population. (c) Fragments Per Kilobase of transcript per Million mapped reads (FPKM) of CNS cell-type specific markers. (d) FPKM values of leukocyte binding adhesion molecules including *Vcam1*. (e) FPKM values of the gene transcripts in murine young and aged CD31+BECs of human plasma proteins that change with age (see **Supplemental Table 2** for list of human plasma proteins expressed in murine BECs) (f) C57BL6 mice were injected with anti-VCAM1-DL488 or IgG isotype control (r.o.) 2 hours before perfusion to label BECs *in vivo* prior to brain dissociation, staining and FACS. (g) Gating and histogram plots of CD31+VCAM1+ cells sorted from the hippocampi of LPS stimulated aged (19-month old) wildtype mice injected with fluorescently tagged anti-VCAM1 mAb or IgG isotype (h) Quantification of CD31+VCAM1+cells isolated from healthy cortex (n=5 single mice/age group). **p<0.002 (i) Quantification of CD31+VCAM1+cells isolated from 4 pooled hippocampi (technical replicates shown). **p<0.005 (j) Flow gating and histogram plots of pooled (n=4 mice/ age group), young or aged hippocampi isolated from healthy mice also injected with anti-VCAM1 mAb. (k) sVCAM1 ELISA in plasma from young isochronic or heterochronic parabionts following 5 weeks of parabiosis. n=11 mice/group pooled from two independent experiments. **p<0.004, Student’s *t-test*. All error bars indicate SEM. (l) Confocal images in the DG of VCAM1, lectin, and Aqp4 of young isochronic or heterochronic parabionts 5 weeks after surgery. Hoechst labels cell nuclei. Scale bar = 100 μm.

**Supplementary Figure 2.**
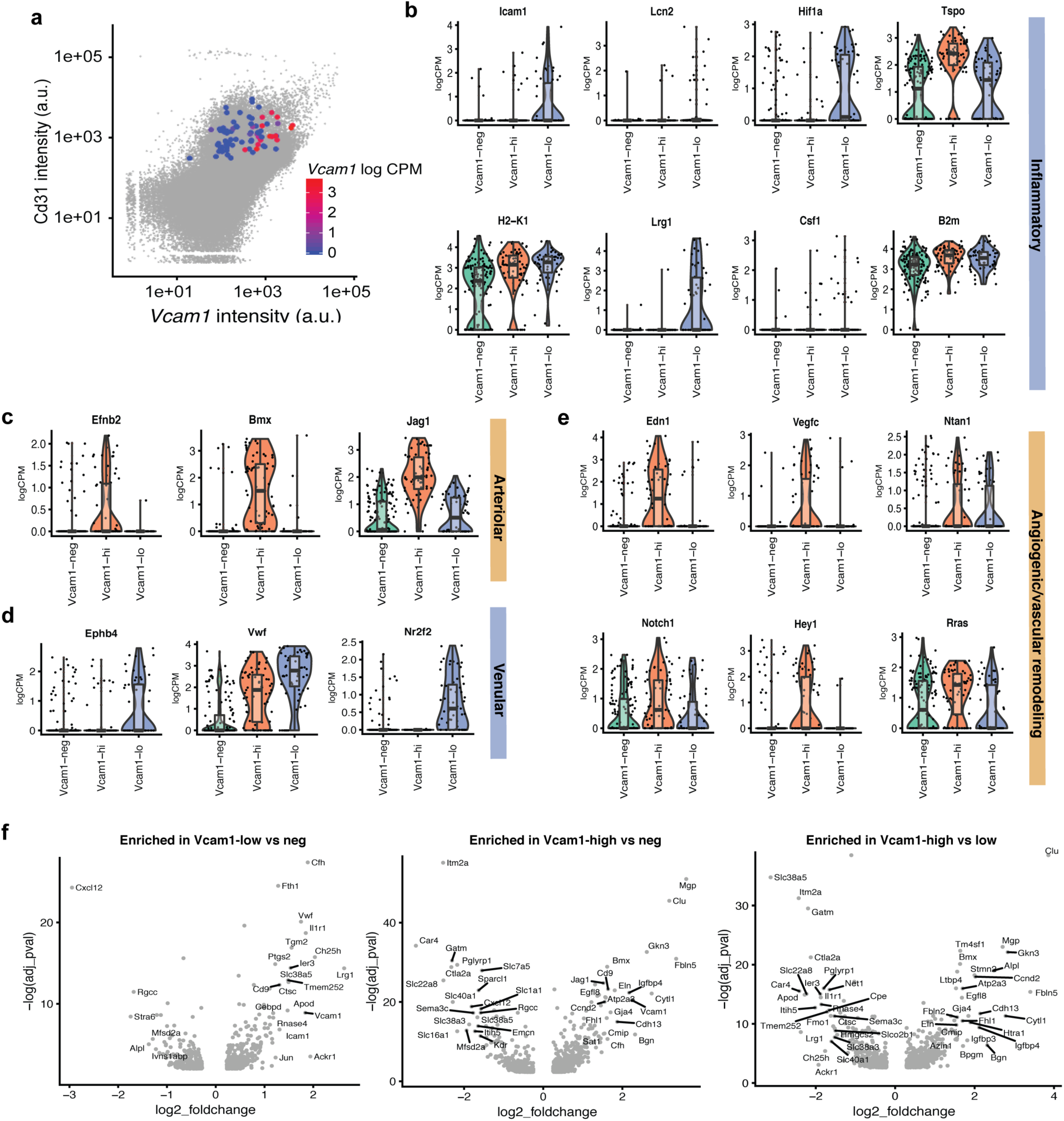
Single Cell Transcriptome profiling of VCAM1 enriched brain endothelial cells reveal specialized subclusters. *Related to Figure 2* (a) Overlay of *Vcam1* mRNA levels on corresponding coordinate on the *Cd31* vs *Vcam1* fluorescent intensity plots obtained during FACs sorting. (b) Violin plots of various inflammation-related genes in each of the 3 distinct clusters. (c) Violin plots of classical arteriolar markers in each cluster. (d) Violin plots of classical venular markers in each cluster. (e) Violin plots of various angiogenesis and Notch-signaling related genes in each of the 3 distinct clusters. (f) Volcano plots of differentially expressed genes when directly compared between Vcam1-high, low and negative clusters. Outlier genes with an adjusted p-value of >0.05 and a log2 fold change of > 1.2 are labeled for visualization.

**Supplementary Figure 3.**
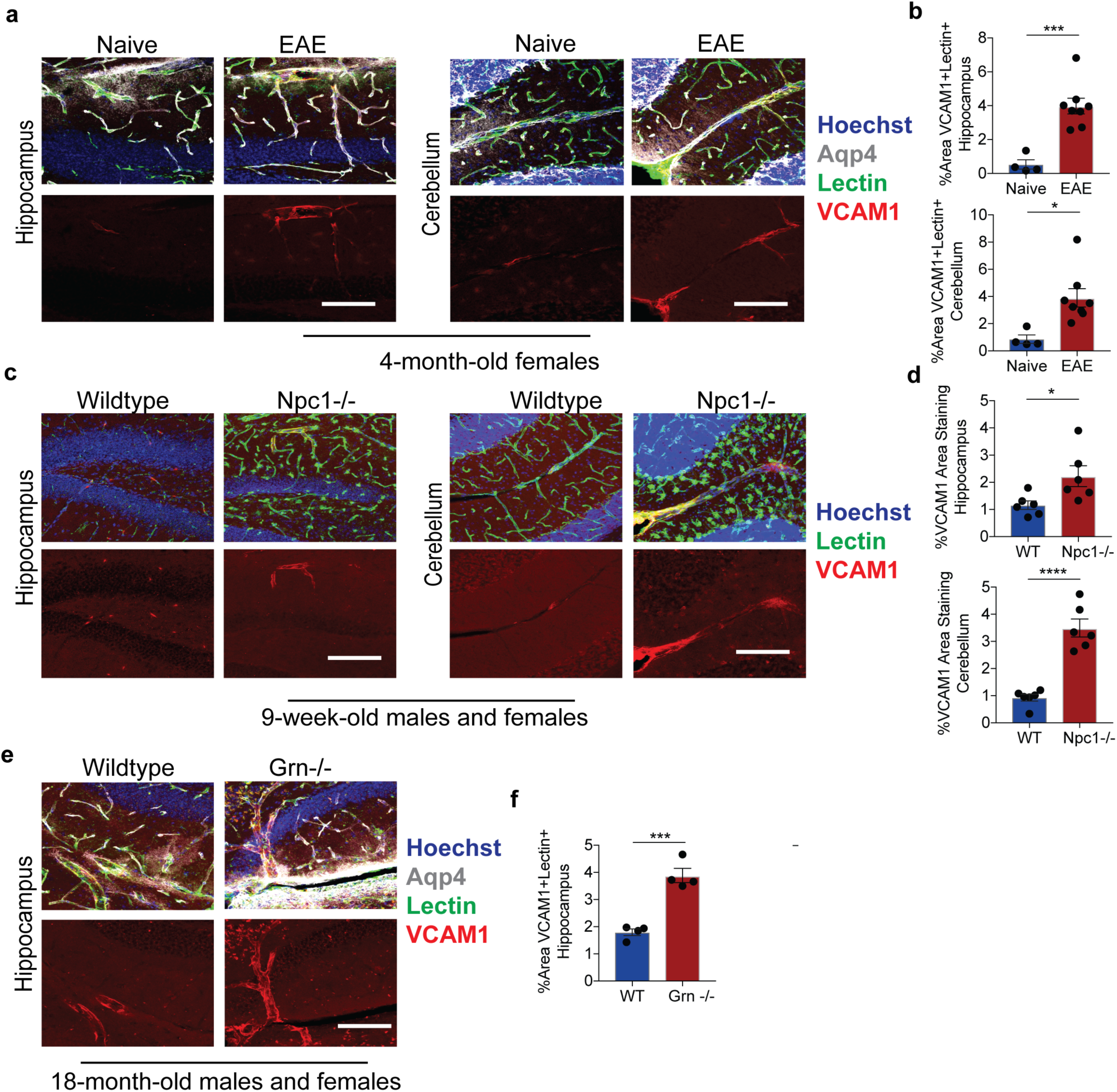
VCAM1 is increased in various pro-inflammatory and neurodegenerative disease models (PGRN, EAE, NPC). *Related to Figure 1* (a) Representative confocal images and quantification (b) of VCAM1, Aqp4, Lectin, with Hoechst labeling cell nuclei in the hippocampus and cerebellum of an EAE (multiple sclerosis) model. Scale bar = 100 μm. ***p<0.0007, *p<0.02. (c) Representative confocal images and quantification (d) of VCAM1, Lectin, with Hoechst labeling cell nuclei in the hippocampus and cerebellum of a Npc1-/- (Niemann Pick Disease Type C) model. Scale bar = 100 μm. ****p<0.0001, *p<0.03. (e) Representative confocal images and quantification (f) of VCAM1, Lectin, Aqp4, with Hoechst labeling cell nuclei in the Grn-/- (Frontotemporal Dementia) model. Scale bar = 100 μm.***p<0.0005. Student’s *t-test*. All error bars indicate SEM.

**Supplementary Figure 4.**
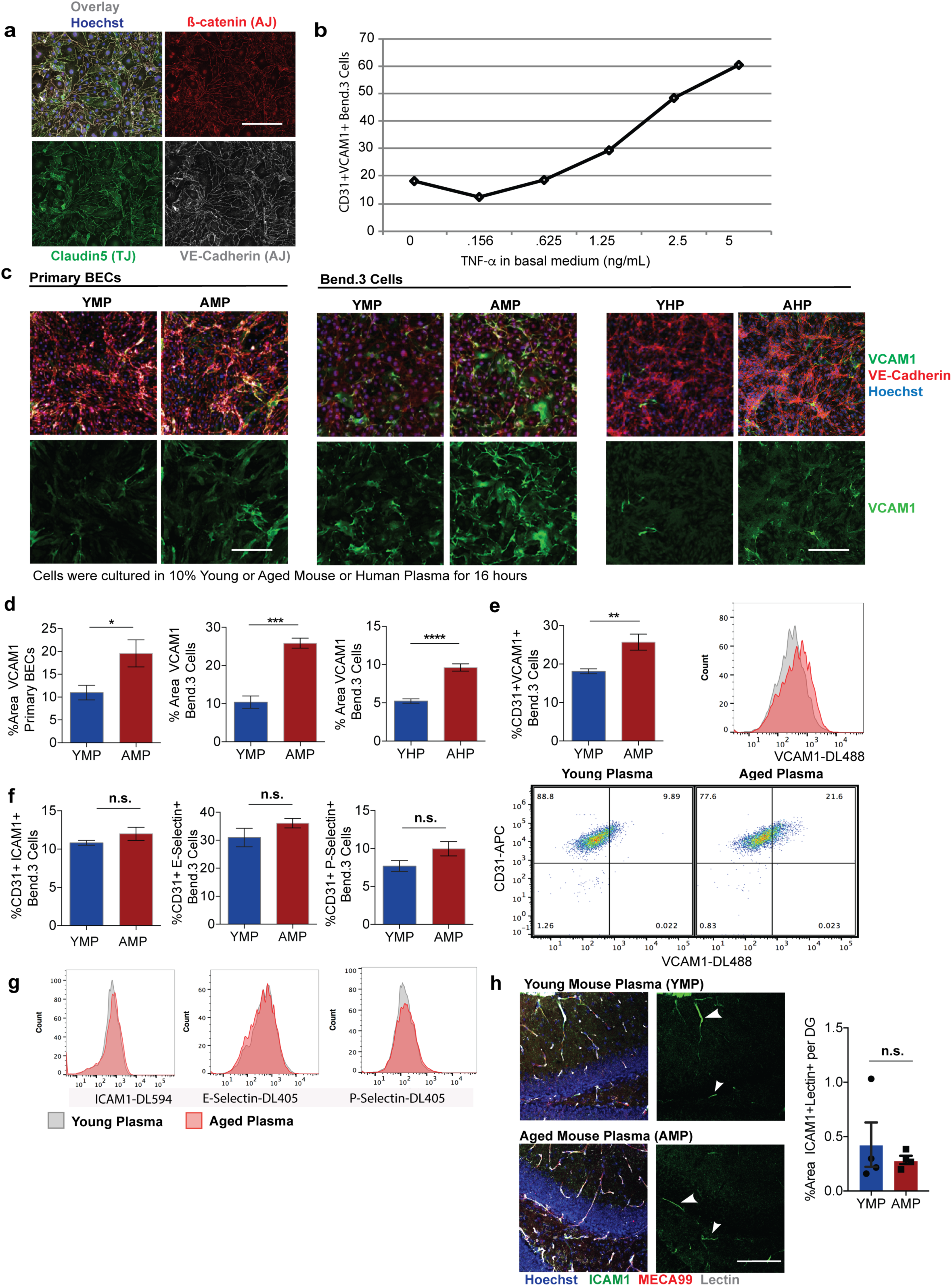
Aged plasma upregulates VCAM1 on cultivated BECs. *Related to Figure 1 and Figure 3* (a) Representative images of Bend.3 cells immunostained for BBB specific markers of adherens junctions (AJ) and tight junctions (TJ), specifically ß-catenin, Claudin-5, and VE-Cadherin. Hoechst labels cell nuclei. Scale bar = 100 μm. (b) Dose response graph depicting cultured Bend.3 cells stimulated overnight with increasing concentrations of recombinant mouse TNF-α followed by flow cytometry to quantify %CD31 + VCAM1+ cells. n=2 pooled samples per condition. (c) Primary BECs and Bend.3 cells cultured in 10% young or aged mouse plasma (YMP: 3-month old; AMP: 18-month-old) or young or aged human plasma (<25 years or >65 years, YHP/AHP) for 16 hours then stained for VE-Cadherin, VCAM1, and Hoechst to label cell nuclei. Representative images are shown. Scale bar = 100 μm. (d) Quantification of VCAM1 %area staining. *p<0.04, ***p<0.0004, ****p<0.0001 n=4-6/group, student’s *t-test*. All error bars indicate SEM. (e) Bend.3 cells cultured in 10% young or aged mouse plasma (YMP/AMP) for 16 hours followed by flow cytometry of CD31 and VCAM1. Graph of %CD31+VCAM1+ quantification shown with histogram and flow cytometry gating of Bend.3 cells. **p<0.005, student’s *t-test*. (f) Quantification of %CD31+ cells colabeled with ICAM1, E-Selectin, or P-Selectin. n=6 replicates/group. All error bars indicate SEM. Histogram plots shown in (g). (h) Representative images of ICAM1, Meca99, lectin, and Hoechst to label cell nuclei of young (3-month-old) mice which received 7 r.o. injections of young (3 mo) or aged (18 mo) pooled plasma over 4 days as described in Figure 2A schematic. Quantification on the right. Scale bar = 100 μm.

**Supplementary Figure 5.**
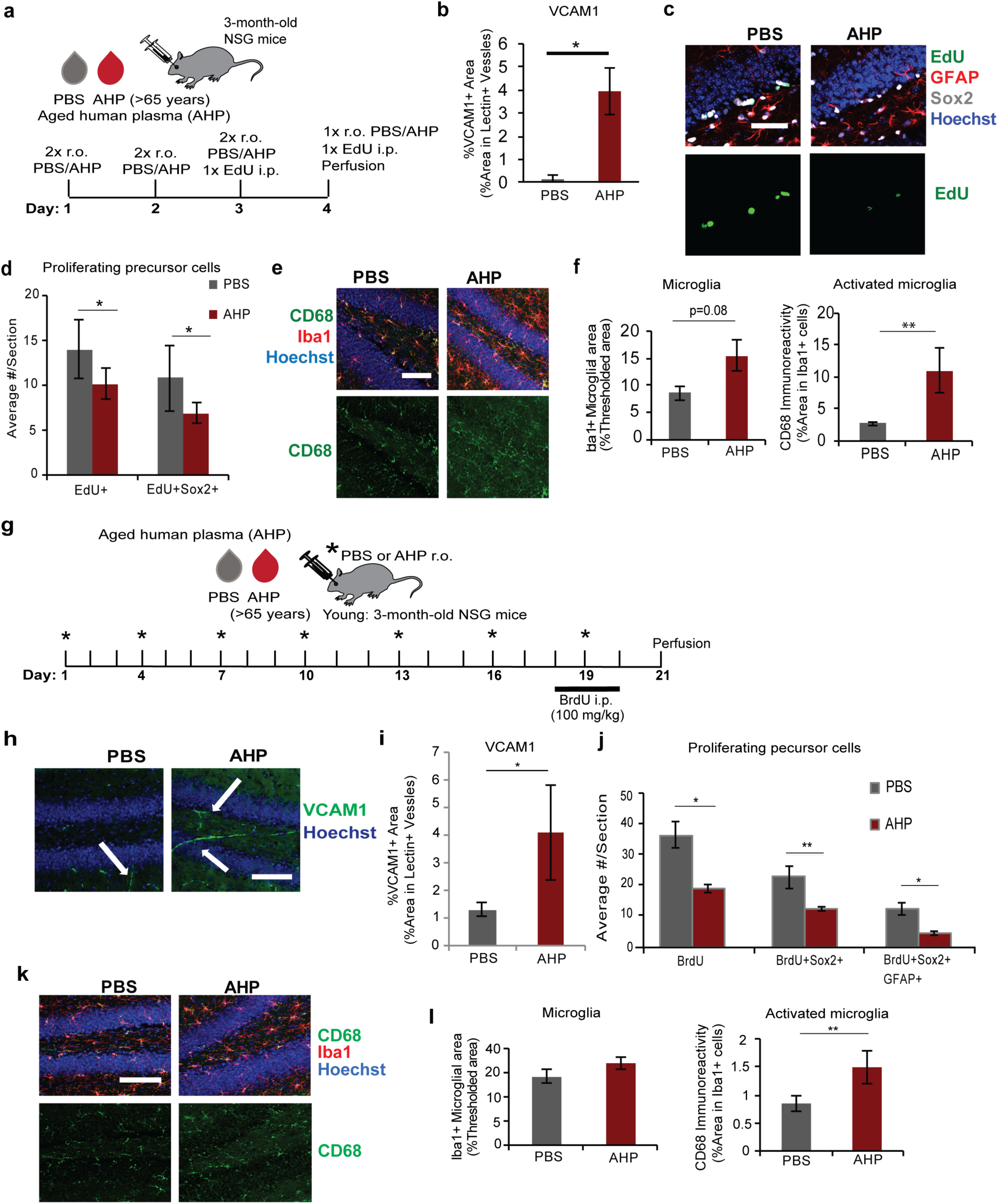
Immunodeficient mice exposed to aged human plasma over 4 days or 3 weeks have increased cerebrovascular VCAM1, decreased neurogenesis, and increased microglial reactivity. *Related to Figure 3* (a) Experimental design. n=5 mice/group (b) Quantification of %VCAM1+Lectin+ staining. *p<0.03. (c) Representative confocal images and quantification (d) in the DG of EdU+ proliferating cells and EdU+ and Sox2+ colabeled proliferating neural progenitor cells. GFAP labels astrocytes and neural stem cells. Scale bar = 50 μm. *p<0.05. (e) Representative confocal images and (f) quantification in the DG of CD68, Iba1 and Hoechst to label cell nuclei. Scale bar = 100 μm. p=0.08; **p<0.05. Student’s *t-test*. All error bars represent SEM. (g) Schematic. n= 6-7 mice/group. (h) Representative confocal images and (i) quantification in the DG of VCAM1 (arrows). Hoechst labels cell nuclei. *p<0.05. Student’s *t-test*. Scale bar = 100 μm. (j) Quantification in the DG of BrdU+ and Sox2+ neural precursor cells and triple labeled GFAP+ neural stem cells from confocal images of immunostained sections. Scale bar = 100 μm. *p<0.02, **p<0.05. (k) Representative confocal images and quantification (l) in the DG of Iba1, CD68, and Hoechst to label cell nuclei. Scale bar = 100 μm. **p<0.05. Student’s *t-test*. Error bars represent SEM.

**Supplementary Figure 6.**
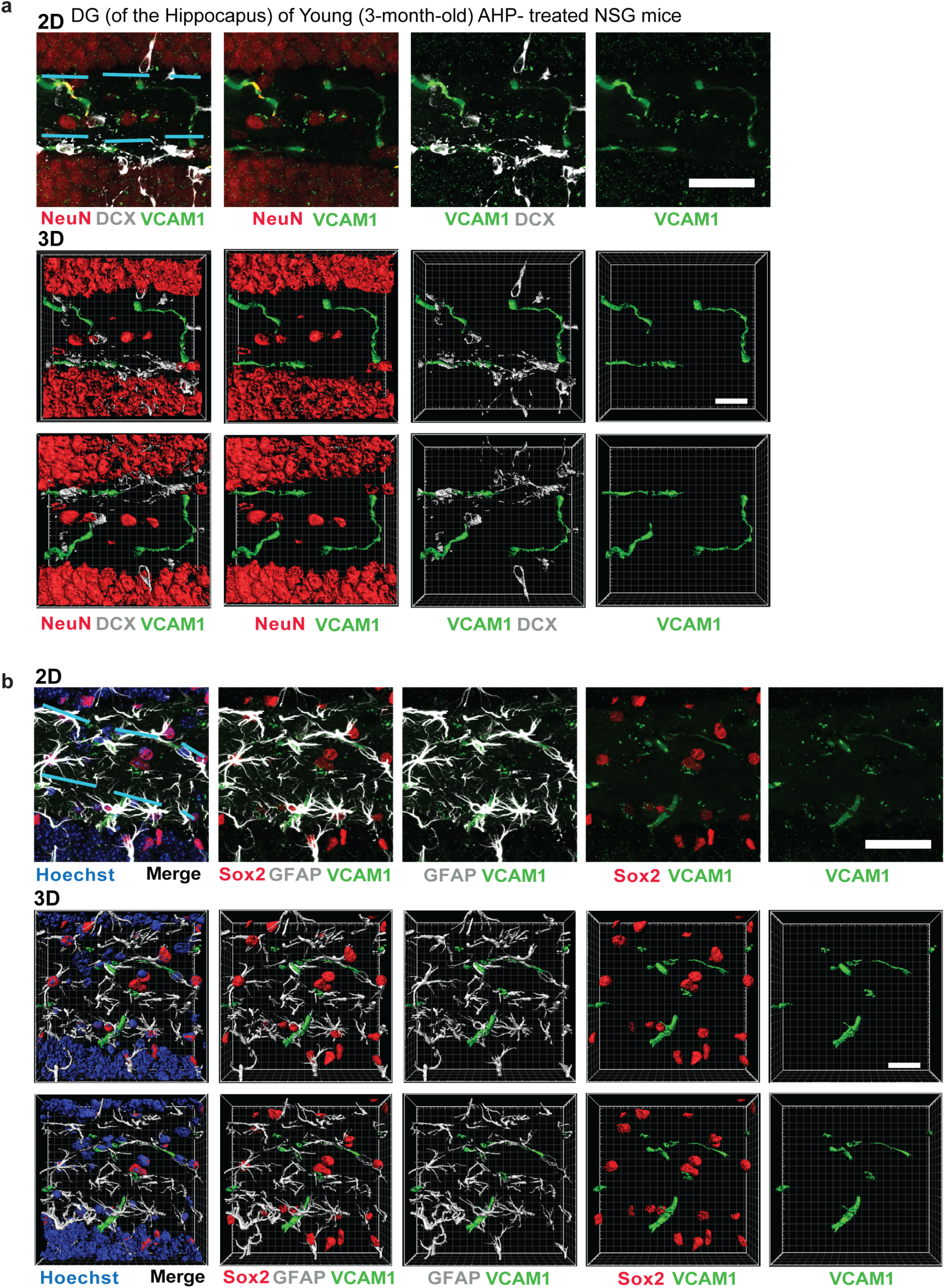
VCAM1 is expressed in Brain Endothelium, but not in other CNS cell types in the hippocampus. *Related to Figure 4* (a) Representative 2D and 3D Z-stacked high magnification confocal images (51 slices with an interval of 0.4 um) of VCAM1 in the granular layer of the DG of the hippocampus of a young (3-month-old) NSG mouse acutely treated with Aged Human Plasma (AHP). Brain sections were co-stained with DCX and NeuN to label immature and mature granule neurons, respectively. VCAM1 is not expressed in these cell types. Light blue lines outline the granule layer. 2D Scale bar = 50 μm. Two 3D renderings of the 2D images are displayed with 180°rotations. 3D Scale bar = 20 μm. (b) Representative 2D and 3D Z-stacked high magnification confocal images (51 slices with an interval of 0.4 um) of VCAM1 in the granular layer of the DG of the hippocampus co-stained with Sox2 and GFAP to label neural stem and progenitor cells (Sox2+GFAP+) and hilur GFAP+ astrocytes. VCAM1 is not expressed in these cell types in the DG. Light blue lines outline the granule layer. 2D Scale bar = 50 μm. Two 3D renderings of the 2D images are displayed with 180°rotation. 3D Scale bar = 20 μm.

**Supplementary Figure 7.**
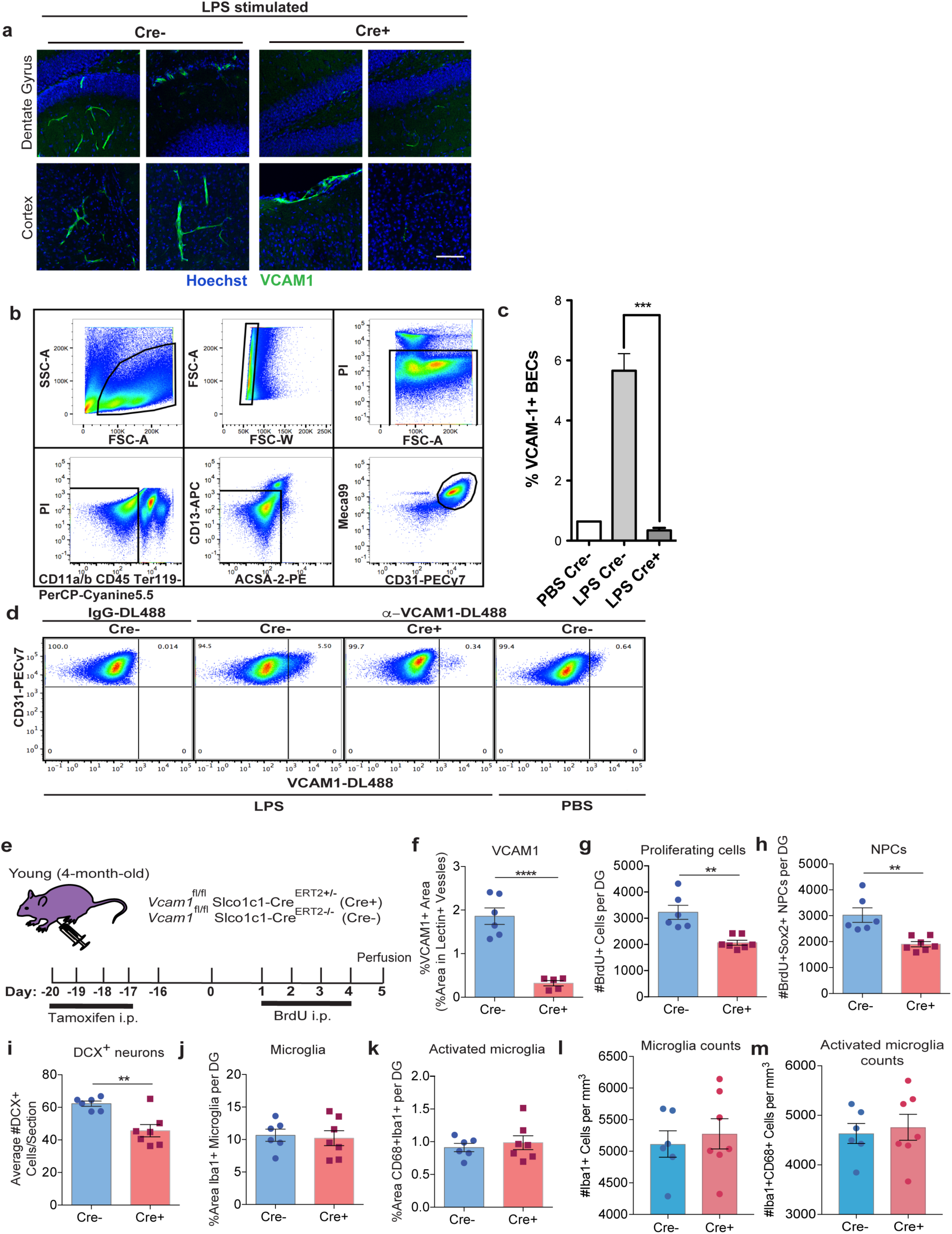
Assessment of Vcam1^fl/fl^Slco1c1-Cre^ERT2+/^' young mice stimulated with LPS and tamoxifen-treated. *Related to Figure 4* (a) Vcam1^fl/fl^Slco1c1-Cre^ERT2+/-^ (Cre+) or Cre^ERT2-/-^ (Cre-) littermates (3-month-old) were treated daily with tamoxifen (i.p. 150 mg/kg) for 5 days followed by 4 days of rest. Mice received 3 LPS injections (0.5 mg/kg i.p.) at 28 hours, 22 hours, and 2 hours prior to perfusion. Mice also received a retro-orbital injection of fluorescently conjugated mouse anti-VCAM1 mAb (100μg) 2 hours prior to perfusion. Representative confocal images of cortex and DG for VCAM1 and Hoechst to label cell nuclei. Loss of Vcam1 in Cre+ mice, but not Cre-, in BBB endothelium, but not in meninges is shown. Scale bar = 100 μm. (b) FACS gating strategy to isolate single BECs. PI+ dead cells were excluded. CD11a/b, CD45, and Ter-119 negative cells were gated to exclude erythrocytes, monocytes/macrophages and microglia. CD13 and ACSA-2 staining was applied to exclude pericytes and astrocytes, respectively. CD31+MECA99+ cells were defined the BEC population. (c) Quantification of (d) flow cytometry that was performed on primary BECs isolated from Cre+ or Cre-mice treated as described in (a). n=3 Cre+ or Cre-mice received LPS, while one Cre-mouse was given PBS vehicle control instead. The VCAM1 gate was set based on a Cre-mice injected with fluorescently conjugated IgG. ***p<0.0007; Unpaired Student’s *t-test*; Error bar represents SEM. (e) Experimental Design. n= 6-7 mice/group. (f) Quantification of VCAM1+ percent area in lectin+ vasculature of immunostained sections from 5–6 mice/group. ****p<0.0007; Student’s *t-test*. (g-i) Quantification of the total number of BrdU+ cells, BrdU+Sox2+ co-labeled neural progenitor cells, and DCX+ immature neurons in the DG of immunostained sections. **p<0.003, Student’s *t-test*. (j-m) Quantification of Iba1 and CD68 in the DG of immunostained sections. All error bars represent SEM.

**Supplementary Figure 8.**
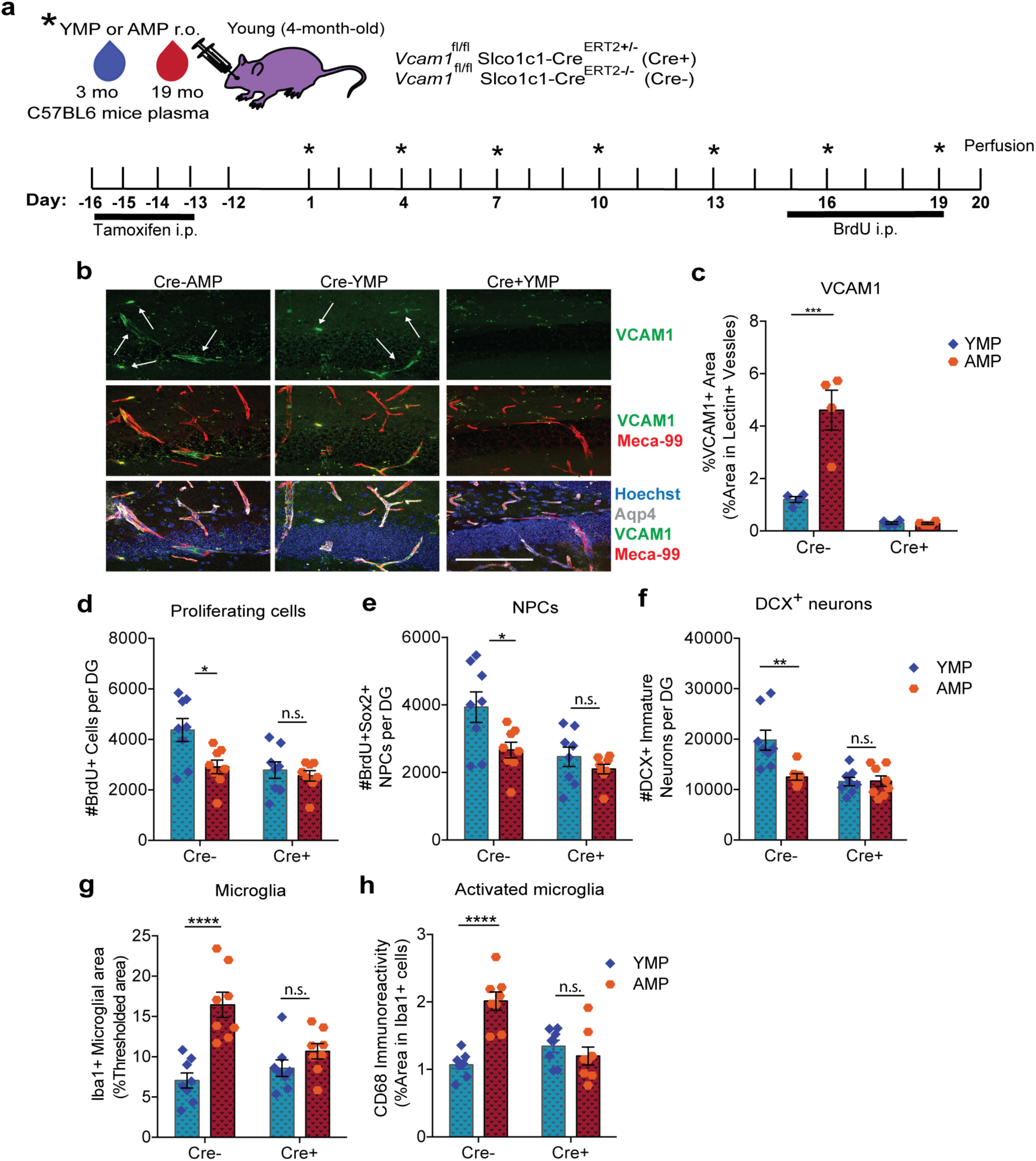
Brain endothelial and epithelial-specific *Vcam1* deletion in young mice mitigates the negative effects of 3 weeks of aged plasma administration. *Related to Figure 4* (a) Experimental Design. n=8 mice/group. (b) Representative confocal images and (c) quantification in the DG of VCAM1, MECA-99, and Aqp4. Hoechst labels cell nuclei. Scale bar = 100 μm. Arrows point to VCAM1+ vessels. (4 mice/group analyzed). ***p<0.0009. (d-f) Quantification of the total number of BrdU+ cells, BrdU+Sox2+ neural progenitor cells, and DCX+ immature neurons in the DG of immunostained sections. *p<0.02, **p<0.007. (g-h) Quantification of Iba1 and CD68 in the DG of immunostained sections. ****p<0.0001, 2-way ANOVA with Tukey’s *post-hoc* test. All error bars represent SEM.

**Supplementary Figure 9.**
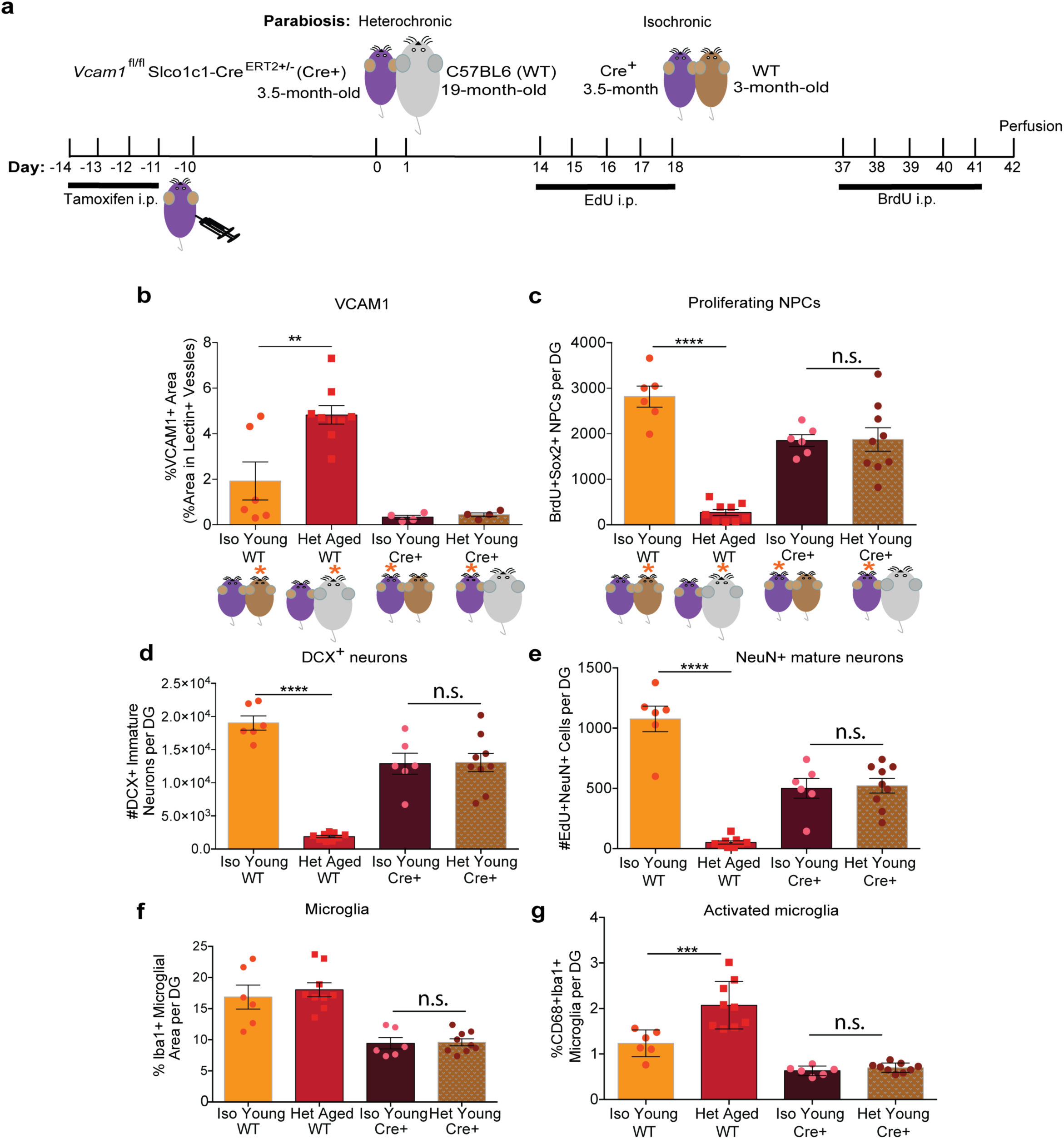
Brain endothelial and epithelial-specific *Vcam1* deletion in young mice mitigates the negative effects of heterochronic parabiosis on young mice. *Related to Figure 4* (a) Experimental Design. n=6 isochronic young pairs (3-month-old C57BL6/J mice and 3.5-month-old Cre+ mice) and 9 heterochronic pairs (18-month-old C57BL6 mice and 3.5-month-old Cre+ mice). (b) Quantification of VCAM1+ percent area in lectin+ vasculature of immunostained sections. **p<0.0001. (c-d) Quantification of the total number of BrdU+Sox2+ neural progenitor cells and DCX+ immature neurons in the DG of immunostained sections. ****p<0.0001. (e) Quantification of the total number of EdU+NeuN+ mature neurons in the DG of immunostained sections. ****p<0.0001. (f-g) Quantification of Iba1 and CD68 in the DG of immunostained sections. ***p<0.0001. 1-way ANOVA with Tukey’s *post-hoc* test. All error bars represent SEM.

**Supplementary Figure 10.**
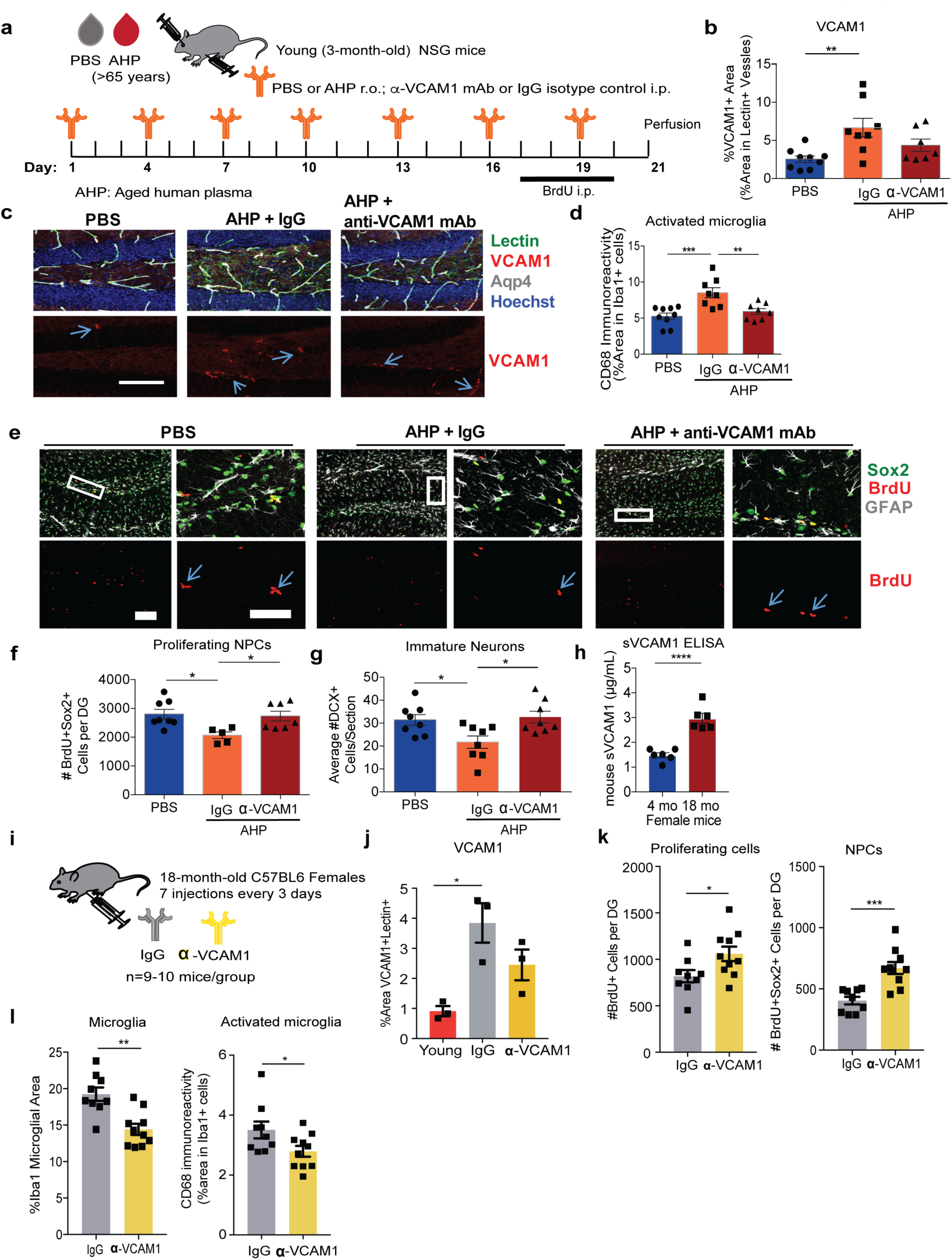
Anti-VCAM1 antibody prevents inhibitory effects of aged human plasma administered over 3 weeks on hippocampal neurogenesis and microglial reactivity and rejuvenates aged female mice brains. *Related to Figures 6-7* (a) Schematic. n=10, 3-month-old NSG mice per group. (b) Quantification and (c) representative confocal images in the DG of VCAM1 (arrows), lectin, and Aqp4. Hoechst labels cell nuclei. Scale bar = 100 μm. **p<0.009. (d) Quantification in the DG of CD68 in Iba1+ stained microglia. ***p<0.0007, 1-way ANOVA with Tukey’s *post-hoc* test. Error bars represent SEM. (e) Representative confocal images in the DG of BrdU+, Sox2+ and GFAP+ neural precursor cells. Boxed areas in the SGZ of images in low magnification (scale bar = 100 μm) are shown in right panels in high magnification (scale bar = 50 μm). Quantification of BrdU+Sox2+ progenitor cells (f) and DCX+ immature neurons (g) from confocal images. *p<0.02. (h) sVCAM1 ELISA of the plasma of young (4-month-old) and aged (19-month-old) female mice. ****p<0.0001. (i) Schematic. Aged (18-month-old) C57BL6/J female mice received i.p. injections of a mouse specific anti-VCAM1 mAb or IgG isotype control (9 mg/kg) every 3 days for a total of 7 injections. Mice also received BrdU daily (100 mg/kg i.p.) for 6 consecutive days followed by perfusion 2 days after the last injection. n=9–10 mice/group. (j) Quantification of VCAM1+Lectin+ staining from confocal images in the DG. *p<0.02, 1-way ANOVA with Tukey’s multiple comparisons *post-hoc* test. (k) Quantification of BrdU+ and BrdU+Sox2+ staining from confocal images in the DG. ***p<0.0004, *p<0.04. (l) Quantification of Iba1 and CD68 staining from confocal images in the DG. **p<0.001, *p<0.05, Student’s *t-test* unless otherwise noted. All error bars represent SEM.

**Supplementary Figure 11.**
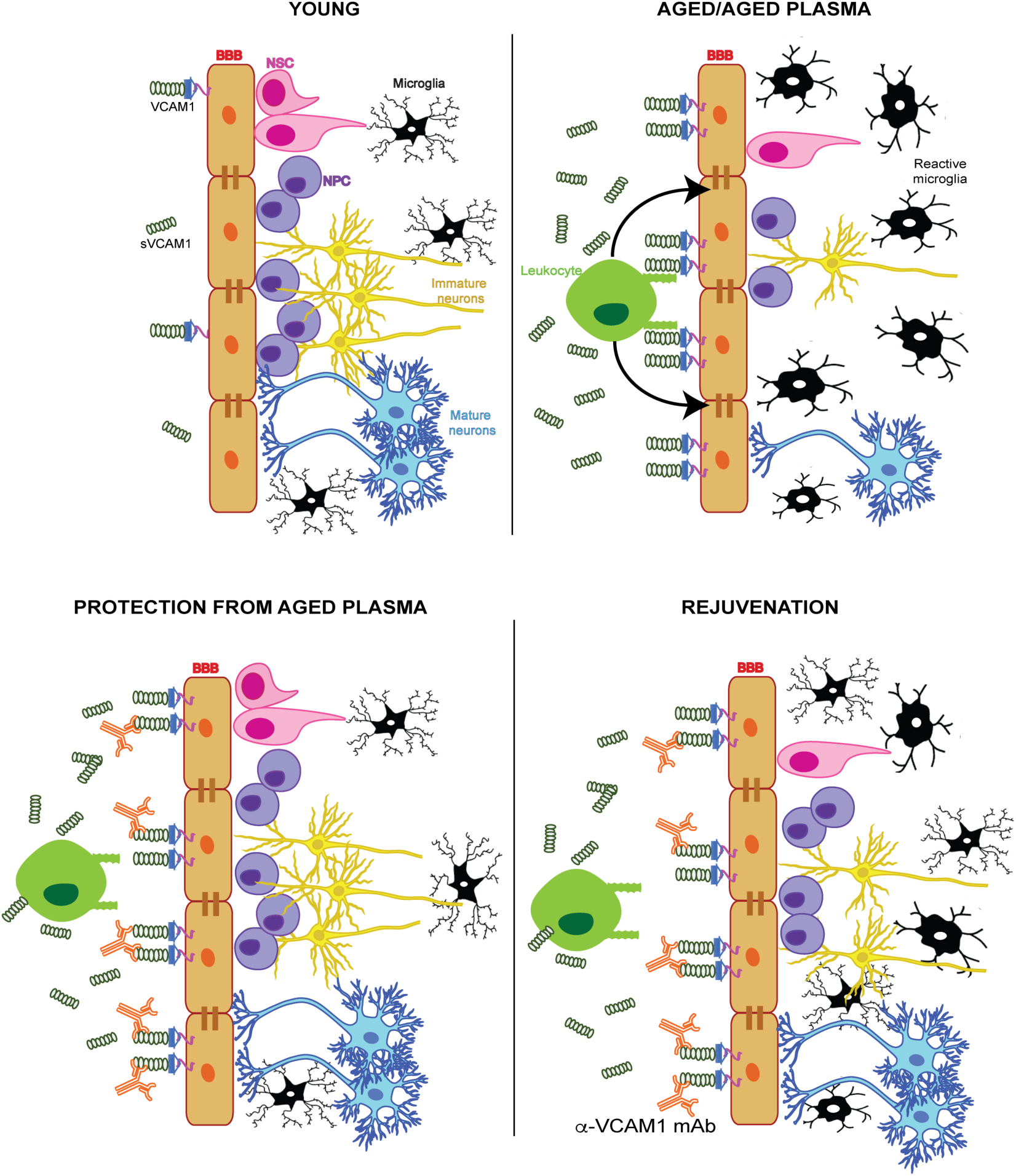
Cartoon Schematic: Aged blood inhibits hippocampal neurogenesis and activates microglia through VCAM1 at the blood-brain barrier. In young healthy mice, neurovascular homeostasis is maintained with low expression levels of systemic and BBB-specific VCAM1, active neurogenesis and nonreactive microglia in a low inflammation environment. During aging or exposure to aged plasma, we propose: 1) Factors in aged plasma induce BEC activation and upregulation of VCAM1. 2) VCAM1 facilitates tethering, but not transmigration, of leukocytes which sustain BEC inflammation. 3) Inflamed VCAM1+ brain endothelium relay (unknown) signals to the parenchyma leading to a loss of homeostasis, decline in neurogenesis and chronic activation of microglia.

## Online Methods

### Human blood samples

Human blood samples from healthy males in the age range of 18–25 and 65–74 were anonymously donated to the Stanford Blood Center, Palo Alto. Blood was centrifuged at 1600g for 10 min at 4°C, plasma was collected and centrifuged again at 1600g for 10 min at 4°C. Plasma was dialyzed using cassettes (Slide-A-Lyzer Dialysis Cassettes, 3.5k MWCO, 12 ml (Fisher, PI 66110)) in 4 L phosphate buffered saline (PBS) with stir bar for 45 min at room temperature, with fresh PBS every 20 min. Cassettes were transferred to fresh 4 L PBS with stir bar for overnight dialysis at 4°C. Plasma samples from 5 aged individuals >65 years old were pooled together for aged human plasma injections or in vitro studies. Plasma samples anonymously donated by 5 young adults <25 years old were pooled together for in vitro studies. Plasma was aliquotted to prevent more than 1 freeze-thaw and stored at −80°C until further use.

### Plasma collection, dialysis and processing

<u>Mouse:</u> Approximately 500 μl of blood was drawn from the heart in 250 mM EDTA (Sigma Aldrich, CAS Number: <u>60-00-4</u>) and immediately transferred to ice. Blood was centrifuged at 1000g for 10 min at 4°C with a break set to 5 or less. Plasma was collected and immediately snap frozen on dry ice and stored at −80°C until further processing. Plasma was dialyzed in 4L of 1X PBS (51226, Lonza) stirred at room temperature. Plasma was transferred to a fresh 4L of 1 X PBS after 45 min and then again 20 min later. After the second transfer, plasma was dialyzed overnight at 4°C in 4 L of stirred 1X PBS. Plasma from 7–9 mice was pooled for injections.

<u>Human:</u> Donor plasma (healthy males, aged 18-25 year or 65–74 years) was purchased from the Stanford Blood Bank. Human plasma was dialyzed as described for mouse plasma (see above). Plasma from 5 individuals of an age group was pooled for injections.

### Proteomics (Human plasma, VCAM1 analysis)

Britschgi et al. measured plasma factors in cognitively normal and AD patients by multiplex assay which measured 74 cytokines, chemokines, growth factors and related proteins in plasma using bead-based multiplex immunoassays as described ^10^. We used the raw plasma data generated in which the low values were replaced with lowest detectable value measured in AD patients or controls, respectively (“Data 4” of the supplemental data) and focused our analysis on subjects cognitively normal (n=118 subjects, 59 males, 59 females). The age range was between 50 and 88 with a median age of 68. We replaced QNS (quantity not sufficient) values in the dataset by NA and log_10_ transformed the data. To measure the strength of the relationships between age and plasma factors levels, we used R Segmented package ^64,65^ to calculate Spearman’s correlation coefficient. To visualize the changes of plasma factors levels with aging, mean value per decade was calculated for each plasma factors and hierarchical clustering applied.

### Animals

NOD-scid IL2Rγ^null^ (NSG) immunodeficient mice were purchased from Jackson Laboratory (Bar Harbor, Maine). NSG mice were bred and only males used for plasma treatment studies. Heterozygous Slco1c1-Cre^ERT2^ breeding males were provided by Professor Markus Schwaninger ^37^. Mice were bred and crossed with Vcam1^fl/fl^ mice (B6.129(C3)-*Vcam1^tm2Flv^*/J mice) purchased from Jackson Laboratory (Bar Harbor, Maine). Male mice were used for plasma treatment studies following treatment with tamoxifen (an estrogen modulator). Aged (greater than 12 months of age) C57BL6J males and females were obtained from the National Institute on Aging (NIA), and young C57BL6J males (2–4 months of age) were purchased from Jackson Laboratory and Charles River. BALB/cNctr-*Npc1^m1N^*/J 9-week-old homozygous males and females were generated by breeding heterozygous mice acquired from Jackson. 17–18-month-old male and female wildtype and Grn-/- deficient mice (B6.129S4(FVB)-*Grn^tm1.1Far^*/Mmja) were bred and aged in-house but originally acquired from Jackson. These transgenic strains were bred and aged in-house and lived under a 12-hour light-dark cycle in pathogenic-free conditions, in accordance with the Guide for Care and Use of Laboratory Animals of the National Institutes of Health.

### In vivo animal studies

#### Parabiosis

Isochronic and heterochronic parabiosis was performed as described ^9^. In brief, mirror-image incisions through the skin were made on the left and right flanks of mice and shorter incisions were made through the abdominal wall. Parabionts were sutured together at their adjacent peritoneal openings. The parabionts’ elbow and knee joints were also sutured together and the skin of each mouse was stapled together. For 1 week during recovery post-surgery, each parabiont mouse received daily subcutaneous injections of Baytril antibiotic solution (5 micrograms per gram of body weight in saline to give a volume of approximately 1% of weight of each mouse) and Buprenorphine (0.1 milligram per milliliter, 0.05 mg/kg) as well as physiological saline (0.9%) for pain relief, prevention of infection and hydration and were monitored regularly.

#### Plasma injections and antibody treatments

**C57BL6J mice:**

Young (3-month-old) C57BL6J male mice were treated with 7 injections of young (3-month) or aged (18-month) dialyzed and pooled mouse plasma (150 uL, r.o.), coming from 8–10 mice per pooled plasma sample. Mice were treated acutely over 4 days, with 2 injections per day spaced 10–12 hours apart. On day 3 during both morning and evening injections and the morning of day 4, mice were pulsed with BrdU to label proliferating cells (100 mg/kg, i.p.; B5002-5G, Sigma Aldrich). Mice received a 7^th^ plasma injection along with BrdU on day 4 followed by perfusion 3 hours after the last injection.

#### Aged mice treated with anti-Vcam1 monoclonal antibody

Aged (16-month-old) C57BL6J mice received i.p. injections of anti-VCAM1 mAb or IgG isotype control (9 mg/kg) every 3 days for a total of 7 injections. Mice also received BrdU daily (100 mg/kg i.p.) for the last 6 days prior to perfusion 24 hours after the last BrdU injection and 48 hours after the last antibody injection.

EAE was induced in young wildtype C57BL6J (4-month-old) female mice as previously reported ^66^.

**NSG mice plasma treatments:**

#### Long-term plasma treatment

NSG mice received PBS or pooled aged human plasma (AHP, >65 years, dialyzed plasma from 5 individuals pooled, 150 uL/injection, r.o) every 3 days for 3 weeks, totaling 7 injections. They also received daily EdU injections (150 mg/kg, i.p.) for 3 days, beginning two weeks after plasma treatment, followed by daily BrdU injections (150 mg/kg, i.p.) beginning on day 18 for 3 days followed by perfusion. n= 6-7 mice/group.

#### Acute Plasma treatment

NSG mice received PBS or pooled aged human plasma (AHP, >65 years, dialyzed plasma from 5 individuals pooled, 150 uL/injection, r.o) twice daily in morning and evenings 10–12 hours apart for 7 total injections. They also received EdU (150 mg/kg, i.p.) 16 hours and 4 hours before perfusion. n=5 mice/group

#### Acute plasma paradigm with anti-VCAMI

3-month-old NSG mice received rat anti-mouse VCAM1 mAb or rat IgG isotype control (9 mg/kg) on day 0 and day 3. Mice were given r.o. injections (150 μl) of aged human plasma (AHP, >65 years, pooled from 5 individuals) or PBS as control twice daily for 7 total injections. Mice were pulsed with EdU (100 mg/kg, i.p.) 16 hours and 2 hours before perfusion to label proliferating cells.

#### Longterm plasma paradigm with anti-VCAM1

NSG mice received pooled aged human plasma (AHP, >65 years) injections (150 uL, r.o.) every 3 days for 7 total injections. In addition, mice received i.p. injections of a anti-VCAM1 blocking mAb or IgG isotype control (9 mg/kg) every 3 days for a total of 7 injections. They also received daily BrdU injections (150 mg/kg, i.p.) for 4 days beginning on day 16.

#### Depletion of sVCAMI

Soluble VCAM1 (sVCAM1) was immunoprecipitated from pooled aged human plasma (65–74 years of age, n=5) using superparamagnetic microbeads conjugated to a mouse anti-human-VCAM1 antibody (BBA5, Novus Biologicals) (or monoclonal mouse anti-human IgG antibody (MAB002, R&D Systems) as a control). In order to first conjugate the dynabeads to the anti-IgG or anti-VCAM1 antibodies, 500 μl of (0.5 mg/ml) antibody was added to 25 mg of dynabeads and incubated on a rotator overnight at 37°C and prepared according to manual instructions (14311D, Thermo Scientific]. The following day, 8 mg of conjugated VCAM1 mAb bound to dynabeads (or equal amount of IgG mAb bound to dynabeads) were added to aliquots of 0.5 mL of dialyzed pooled human plasma and incubated at 4C rotating overnight. Depleted plasma was collected the next day and magnetic VCAM1 saturated dynabeads were removed using a magnetic bar through serial transfers of the plasma to new tubes, and stored at −80°C until mouse treatment.

#### NSG Mouse VCAM1-depleted plasma treatment

Following pull-down of sVCAM1, pooled, depleted aged human plasma (IgG versus sVCAM1 depleted) or saline (200 μl/mouse) was injected retro-orbitally (r.o.) into young (4-month-old) NSG mice (n=7-8 mice/group) twice daily for 4 days for 7 total injections. Mice were also injected with BrdU (100 mg/kg, i.p.) starting on the third day. Mice were anesthetized with avertin followed by saline perfusion on the 4^th^ day of treatment, 4 hours after the 3rd BrdU and 7^th^ plasma injections. One mouse per group received intra-orbital injections of 100pg fluorescently labeled (DyLight™488, Thermo Scientific, 53025) InVivoMAb anti-mouse CD106 (VCAM-1, clone M/K-2.7, Bioxell, BE0027) and fluorescently labeled (DyLight^TM^550, Thermo Scientific, 84530) rat anti-mouse MECA-99 (gift of the Butcher lab) 3 hours prior to perfusion.

Slc01c1^CreERT2+/-^; VCAM1^fl/fl^ (Cre+) or Slc01c1^CreERT2-/-^; Vcam1^fl/fl^ (Cre-) mice

##### Basal neurogenesis

Young (4-month-old) heterozygous *Vcam1*^fl/fl^ Slco1c1-Cre^ERT2+/-^ mice (Cre+) or Cre^ERT2-/-^ (Cre-) littermates were injected once daily with tamoxifen (Sigma, T5648, prepared in sunflower seed oil (20 mg/mL solution; 100 mg/kg i.p.) for 4 consecutive days, followed by 2 weeks of rest. Mice received BrdU (100 mg/kg, i.p.) once daily for 4 days. Mice were anesthetized with avertin (tribromoethanol; 0.018 ml (2.5%) per gram of body weight), followed by saline perfusion 24 hours after the final BrdU injections. n= 6-7 mice/group.

<u>Acute paradigm:</u> Young (4-month-old) heterozygous *Vcam1*^fl/fl^ Slco1c1-Cre^ERT2+/-^ mice (Cre+) and littermates controls lacking a copy of Cre^ERT2^ (Cre-) were injected once daily with tamoxifen (100 kg/mg), prepared in sunflower seed oil (20 mg/mL solution) intraperitoneally (i.p.) for 4 consecutive days. After a 3-day resting period, mice were treated acutely over 4 days with young plasma (3-month-old C57BL6J pooled male plasma, 8–10 mice pooled per sample) or aged plasma (18-month-old C57BL6J pooled mouse plasma, 8–10 mice pooled per sample) (150 uL, r.o.), with 2 injections per day spaced 10–12 hours apart. On day 3 during both morning and evening injections and the morning of day 4, mice were pulsed with BrdU (100 mg/kg, i.p.) to label proliferating cells. Mice received a 7^th^ plasma injection along with a 3^rd^ BrdU injection on day 4 followed by perfusion 3 hours after the last injection. One mouse per group received a single retro-orbital injection of 100μg fluorescently labeled (DyLight™488, Thermo Scientific, 53025) InVivoMAb anti-mouse CD106 (VCAM-1, clone M/K-2.7, Bioxell, BE0027) and fluorescently labeled (DyLight^TM^550, Thermo Scientific, 84530) rat anti-mouse MECA-99 (gift of the Butcher lab) 3 hours prior to perfusion.

<u>Long term paradigm:</u> Young (4-month-old) heterozygous *Vcam1*^fl/fl^ Slco1c1-Cre^ERT2+/-^ mice (Cre+) or Cre ^ERT2-/-^ (Cre-) littermates were injected with tamoxifen (100 mg/kg i.p.) for 4 consecutive days, followed by 12 days of rest. Young or aged pooled mouse plasma (YMP/AMP) (150 uL, r.o.) was injected once every 3 days for 7 total injections. Mice received BrdU (100 mg/kg, i.p.) for 5 consecutive days beginning on 15^th^ day after the start of plasma treatment. Mice were perfused 24 hours after the last BrdU and plasma injection. n=8 mice/group for 4 groups. One mouse per group received r.o. injections of 100μg fluorescently labeled (DyLight™488, Thermo Scientific, 53025) rat anti-mouse VCAM-1 (clone M/K-2.7, Bioxell, BE0027) and fluorescently labeled (DyLight^TM^550, Thermo Scientific, 84530) rat anti-mouse MECA-99 (gift of the Butcher lab) 3 hours prior to perfusion. Mice were anesthetized with avertin followed by saline perfusion 16 hours after the final BrdU injections.

<u>Parabiosis:</u> Young (3.5-month-old) heterozygous *Vcam1*^fl/fl^ Slco1c1-Cre^ERT2+/-^ mice (Cre+) were injected with tamoxifen (100 mg/kg i.p.) for 4 consecutive days, followed by 10 days of rest. Mice then underwent isochronic or heterochronic parabiosis with C57BL6J young (3-month-old) or aged (18-month-old) wildtype mice for 6 weeks. Parabiosis surgery was performed following the Wyss-Coray lab’s previously described protocol ^9^. Briefly, mirror-image incisions through the skin were made on the left and right flanks of mice and shorter incisions were made through the abdominal wall. Parabionts were sutured together at their adjacent peritoneal openings. The parabionts’ elbow and knee joints were also sutured together and the skin of each mouse was stapled together. For 1 week during recovery post-surgery, each parabiont mouse received daily subcutaneous injections of Baytril antibiotic solution (5 micrograms per gram of body weight in saline to give a volume of approximately 1% of weight of each mouse) and Buprenorphine (0.1 milligram per milliliter, 0.05 mg/kg) as well as physiological saline (0.9%) for pain relief, prevention of infection and hydration and were monitored regularly.

Each parabiont mouse was administered EdU (100 mg/kg, i.p.) for 5 days, 2 weeks after parabiosis surgery. Mice received BrdU (100 mg/kg, i.p.) for 5 consecutive days beginning on day 37 after parabiosis surgery. Mice were perfused 24 hours after the last BrdU injection, after 6 weeks of parabiosis. n=6 isochronic young pairs (3-month-old C57BL6J mice and 3.5-month-old Cre+ mice) and 9 heterochronic pairs (18-month-old C57BL6J mice and 3.5-month-old Cre+ mice).

#### Anti-VCAM1 antibody in vivo retro-orbital injections to label CD31+VCAM1+ BECs

**C57BL6J Mice:**

For Gating of VCAM1+ cells: Mice were injected with LPS derived from *Salmonella enterica* Serotype Typhimurium (Sigma, L6511), i.p. 1 mg LPS/kg body weight at three successive time points: 0h, 6h, and 24h ^67^. Control mice were injected with bodyweight corresponding volumes of PBS. Experimental mice received i.p. and s.c. injections of sterile 0.9% saline with 5% glucose to ensure hydration and stable glucose levels during the procedure. Mice received a the third LPS injection followed by retro-orbital injections of either 100μg fluorescently labeled (DyLight™488, Thermo Scientific, 53025) InVivoMAb anti-mouse CD106 (VCAM-1, clone M/K-2.7, Bioxell, BE0027) or fluorescently labeled Rat IgG 1 Isotype antibody (Clone HRPN, Bioxell, BE0088). Two hours after the last LPS injection (26h) mouse brains were harvested for BEC isolation and flow analysis.

Healthy young (3-month-old), aged (19-month-old), or plasma injected (r.o.) young mice were similarly injected (r.o.) with fluorescently labeled anti-VCAM1 mAb and gated for flow cytometry analysis of CD31+VCAM1+ cells from cortex and hippocampi. Gates are based on positive LPS-stimulated mice injected (r.o.) with anti-VCAM1 or IgG control.

**Slc01c1^CreERT2+/-^; Vcam1^fl/fl^ mice:**

Young (4-month-old) *Vcam1*^fl/fl^ Slco1c1-Cre^ERT2+/-^ mice (Cre+) received tamoxifen (100 mg/kg; i.p.) once daily for 5 days. After a 3-day resting period, mice were treated with LPS at 0, 2, and 24 hour time points (0.5 mg/kg, i.p.) and fluorescently labeled anti-VCAM1 mAb (100μg, r.o.) 2 hours prior to cell isolation and flow cytometry analysis.

##### Primary BEC isolation for bulk and single cell RNA-seq, flow analysis following LPS stimulation, and in vitro cultivation

*Primary BEC isolation and quantification of CD31+VCAM1+ cells:*

BEC isolation was based on a previously described procedure ^68^. Briefly, mice were anesthetized with avertin and perfused following blood collection. After thoroughly dissecting the meninges, cortices and hippocampi were collected, minced and digested using the neural dissociation kit according to kit instructions (Miltenyi, 130-092-628). Brain homogenates were filtered through 35 μm in HBSS and centrifuged pellets were resuspended in 0.9 M sucrose in HBSS followed by centrifugation for 15 min at 850xg at 4°C in order to separate the myelin. This step was repeated for better myelin removal.

Cell pellets were eluted in FACs buffer (0.5% BSA in PBS with 2mM EDTA) and blocked for ten min with Fc preblock (CD16/CD32, BD 553141), followed by 20 minute staining with anti-CD31-APC (1:100, BD 551262), anti-CD45-FITC or anti-CD45-APC/Cy7 (1:100, BD Pharmingen Clone 30-F11 553080; Biolegend, 103116), and anti-Cd11b-BV421 (1:100, Biolegend Clone M1/70 101236). Dead cells were excluded by staining with propidium iodide solution (1:3000, Sigma, P4864). Flow cytometry data and cell sorting were acquired on an ARIA II (BD Biosciences) with FACSDiva software (BD Biosciences). FlowJo software was used for further analysis and depiction of the gating strategy. Gates are indicated by framed areas. Cells were gated on forward (FSC = size) and sideward scatter (SSC = internal structure). FSC-A and FSC-W blotting was used to discriminate single cells from cell doublets/aggregates. PI+ dead cells were excluded. CD11b+ and CD45+ cells were gated to exclude monocytes/macrophages and microglia. CD31+Cd11b-CD45-cells were defined as the BEC population and were sorted directly into RNAlater (Life Technologies, AM7020) and stored at −80°C until further processing. If mice were injected with fluorescently labeled anti-mouse VCAM1-DyLight™488 as described above, CD45 was stained in the APC/Cy7 channel, and CD31+VCAM1+ cells were also gated in the APC and FITC channels.

#### RNA Sequencing

##### Bulk RNAseq

Mice brain hippocampi and cortex (2 pooled brains per sample; n=6 young (3-month-old C57BL6/J males) samples or n=6 aged (19-month-old C57BL6/J males) samples were dissected using the neural dissociation kit (Miltenyi, 130-092-628) following perfusion. BECs (average 81,000 cells CD31+CD45-Cd11b-cells per pooled sample) were isolated using multi-channel flow cytometry and sorted directly into RNAlater as described above. Frozen cells were thawed to room temperature before 10 min centrifugation at 1000g. Total RNA was isolated from the cell pellets using the RNeasy Plus Micro kit (Qiagen, 74034). RNA quantities and RNA quality was assessed using the Agilent 2100 Bioanalyzer (Agilent Technologies). All samples passed a quality control threshold (RIN ≥ 8.5) to proceed to library preparations and RNAseq.

Total mRNA was transcribed into full length c-DNA using the SMART-Seq v4 Ultra Low Input RNA kit from Clontech according to the manufacturer’s instructions. Samples were validated using the Agilent 2100 Bioanalyzer and Agilent High Sensitivity DNA kit. 150 pg of full length c-DNA was processed with the Nextera XT kit from Illumina for library preparation according to the manufacturers protocol. Library quality was verified with the Agilent 2100 Bioanalyzer and the Agilent High Sensitivity DNA kit. Sequencing was carried out with Illumina HiSeq 2000, paired end, 2×100 bp depth sequencer. STAR was used to align the transcript reads to the mouse genome, and CuffDiff package was used to generate Fragments Per Kilobase of Transcript Per Million Reads Mapped (FPKM) reads and do differential expression analysis. R was used to do cluster analysis and generate heat maps displaying up or down-differentially regulated genes in aged versus young BECs. GeneAnalytics and GeneCards-packages offered by Gene Set Enrichment Analysis (GSEA) tool was used for GO pathway analysis and classification of highly expressed and differentially expressed genes and pathways in young and aged BECs.

##### Single Cell RNAseq of VCAM1 enriched BECs

4 young (3-month-old) or 4 aged (19-month-old) C57BL6/J males were injected (r.o.) with fluorescently labeled anti-VCAM1 mAb 2 hours prior to sacrifice and gated for single cell isolation of CD31+VCAM1+ cells from pooled hippocampi following perfusion. Gates are based on positive LPS-stimulated mice injected with anti-VCAM1 mAb or IgG control antibody.

Four hippocampi (from both hemispheres) were pooled together from 4 young (3-month-old) or 4 aged (19-month-old) C57BL6/J males and sorted into lysis buffer in 96-well plates then snap frozen and stored at −80 degrees Celsius until RNA extraction and library preparation. Two, 96-well plates per group contained BECs that were 50% enriched for VCAM1 high expression based on flow cytometry gating; unbiased CD31+ cells were also collected into two, 96-well plates per group.

##### cDNA synthesis, library preparation and sequencing

Cell lysis, and cDNA synthesis was performed using the Smart-seq-2 protocol as described previously ^69,70^, with some modifications. After cDNA amplification (23 cycles), cDNA concentrations were determined via capillary electrophoresis and cells were cherry-picked to improve quality and cost of sequencing. Cell selection was done through custom scripts and simultaneously normalizes cDNA concentrations to ~0.2 ng/uL per sample, using the TPPLabtech Mosquito HTS and Mantis (Formulatrix) robotic platforms. Libraries were prepared using the Illumina Nextera XT kits following the manufacturer’s instructions. Libraries were then sequenced on the Nextseq (Illumina) using 2 × 75bp paired-end reads and 2 × 8bp index reads with a 200 cycle kit (Illumina, 20012861) and pooled using the Mosquito liquid handler. Library quality was assessed via capillary electrophoresis on a Fragment Analyzer (AATI) and quantified by qPCR. Samples were sequenced at an average of 700,000 reads per cell.

##### Bioinformatics pipeline

Sequences from the Nextseq were demultiplexed using bcl2fastq, and reads were aligned to the mm10 genome augmented with ERCC sequences, using STAR version 2.5.2b. Gene counts were made using HTSEQ version 0.6.1 p1. We applied standard algorithms for cell filtration, feature selection, and dimensional reduction. First, genes appearing in fewer than 3 cells, cells with fewer than 300 genes, and cells with les than 50 000 reads were excluded from the analysis. Out of these cells, those with more than 30% of reads as ERCC, and more than 5% mitochondrial or 3% ribosomal were also excluded. Counts were log-normalized (log(1+counts per N)), then scaled via linear regression against the number of reads, the percent mapping to ribosomal genes, and percent mapping to mitochondrial genes. To select for relevant features, genes were first filtered to a set of 3000 with the highest positive and negative pairwise correlations. Genes were then projected into low dimensional principal component space using the robust principal component analysis (rPCA). Single cell PC scores and genes loads for the first 20 PCs were analyzed using the Seurat package in R. Briefly, a shared-nearest-neighbor graph was constructed based on the Euclidean distance metric in PC space, and cells were clustered using the Louvain method. Cells and clusters were then visualized using 3-D t-distributed Stochastic Neighbor embedding on the same distance metric. Differential gene expression analysis was done by applying the Mann-Whitney U-test of the BEC clusters obtained using unsupervised clustering. P-values were adjusted via the false discovery rate (FDR) or Bonferroni. All graphs and analyses were generated and performed in R.

GeneAnalytics and GeneCards-packages offered by Gene Set Enrichment Analysis (GSEA) tool was used for GO pathway analysis and classification of enriched genes in each subpopulation.

#### For LPS-treated Slco1c1^CreERT2+/-^; Vcam1^fl/fl^ mice

BEC isolation was based on a previously described procedure ^71^. Briefly, mice were sacrificed using carbon dioxide asphyxiation followed by cervical dislocation. Mouse brains were carefully removed from the scull and stored on ice in Buffer A (153mM NaCl, 5.6mM KCl, 1.7mM CaCl2, 1. 2mM MgCl2, 15mM HEPES; 10mg/ml bovine serum albumin (BSA) fraction V). After thoroughly dissecting the meninges, cortices and hippocampi were collected and washed several times in Buffer A before the tissues was minced and centrifuged at 300g for 7min at 4°C. The pellet was digested in a 1:1:1 volume mix of tissue, Buffer A, and 0.75% collagenase II (Millipore, C2-22) at 37°C for 50min. The tissue was homogenized by thorough shaking after 25 and 50min of digestion and repetitive up and down pipetting of the cell solution at the end of digestion. The enzymatic reaction was stopped by adding Buffer A. After centrifugation (300g, 7 min, 4°C) the pellet was carefully resuspended in PBS containing 25% BSA (Fisher Scientific, BP1600-1) and centrifuged at 1000g, 30min at 4°C in order to separate the myelin and to enrich for capillary fragments. To deplete for red blood cells the pellet was incubated in Red Blood Cell Lysis Buffer (Sigma, R7757) for 1.5min at room temperature with occasional shaking, followed by a wash step in buffer A and centrifugation (300g, 7 min, 4°C). For the second digestion, the pellet was resuspended in buffer A containing 1mg/ml Collagenase/Dispase (Roche, 11097113001) and the mixture was incubated at 37°C for 13min. 1|jg/ml DNase I (Sigma, 10104159001) was added for 2 additional minutes. To quench the reaction buffer A was added and cells were centrifuged at 300g for 7min at 4°C.

For flow cytometry, the enriched BECs were labeled by standard protocols with fluorochrome-conjugated antibodies (identified in antibodies section) in HBSS (Thermo Fisher) containing 10% FBS for 30min on ice. Dead cells were excluded by staining with propidium iodide solution (1:3000, Sigma, P4864). Background fluorescence was determined by the ‘fluorescence-minus-one’ method and for VCAM1 a specific IgG1 Isotype control antibody was used. Flow cytometry data were acquired on an ARIA II (BD Biosciences) with FACSDiva software (BD Biosciences). FlowJo software (TreeStar) was used for further analysis.

Cells were gated on forward (FSC = size) and sideward scatter (SSC = internal structure). FSC-A and FSC-W blotting was used to discriminate single cells from cell doublets/aggregates. PI uptake indicated dead cells, which were excluded. CD11a/b, CD45, and, Ter-119 negative cells were gated to exclude erythrocytes, monocytes/macrophages and microglia. CD13 and ACSA-2 staining was applied to exclude pericytes and astrocytes, respectively. CD31+MECA99+ cells were defined as the BEC population.

BEC cultivation was based on a previously described procedure ^72^. For primary BEC cultivation, cells were resuspended in endothelial cell growth medium (20% FBS, 2mM L-glutamine, 2mM penicillin-streptomycin, 1x MEM non-essential amino acids, 0.1mg/ml heparin, 1mM sodium pyruvate, 1mg/ml sodium hydrogen carbonate, 0.05mg/ml ECGS in MCDB-131) and seeded on 1mg/ml rat tail collagen (BD Biosciences, 354236) coated tissue culture plates. After 24h, 4μg/ml puromycin (Santa Cruz, sc-108071A) was added for 48h to remove potentially contaminating cells ^73^.

All animal experimental procedures were performed in accordance with the Guide for Care and Use of Laboratory Animals of the National Institutes of Health

#### Tissue processing

Mice were anesthetized with Avertin (2,2,2, Tribromoethanol: T48402, Sigma Aldrich; 2-methyl-2-butanol: 240486, Sigma Aldrich) (0.018 ml (2.5%) per gram of body weight) prior to sacrifice. 0.5 ml blood was collected from the right heart ventricle in 250 mM EDTA and kept on ice until further processing. Mice were perfused with 20 ml PBS. Blood was centrifuged at 1000g for 10 min at 4°C, plasma was rapidly frozen with dry ice and stored at −80°C until further processing. Brains were harvested and divided sagittal. One hemisphere was used to dissect hippocampus and cortex which was snap frozen and stored at −80°C. The second hemisphere was fixed in phosphate-buffered 4% paraformaldehyde (PFA) for 72h at 4 °C before transferring to 30% sucrose in PBS at 4 °C for 48 hours. Brains were frozen at −30°C and cryosectioned coronally at 40μm using a microtome (Leica SM2010R). Brain sections were stored in cryoprotectant (40% PBS, 30% glycerol, 30% ethynele glycol) and kept at −20°C until staining.

#### Cell Culture

For all studies, Bend.3 cells (gift of the Butcher Lab; purchased from America Type Culture Collection) were used. Bend.3 cells are immortalized brain endothelial cells isolated from BALB/C mice (CRL-2299, ATCC) ^34^. These cells were seeded at 40,000 cells/cm^2^ in MCDB 131 HUVEC medium (10372019, Life Technologies) supplemented with the following: 15% endotoxin-free fetal bovine serum (SH30071, GE Healthcare Life Sciences), 1% sodium pyruvate (11360070, Life Technologies), 1% heparin (H4784, Sigma Aldrich), 1% pen-strep (1786396, Life Technologies), 1% non-essential amino acids (11140050, Life Technologies), 1% l-glutamine (2 mM, Fisher Scientific, glutamax supplement, 35050061), and 1% 100 mg/mL sodium bicarbonate (S5761, Sigma Aldrich). To confirm cell morphology and tight- and adherens-junctions we stained with B-catenin (Millipore, 05-665), Claudin-5 (Thermofisher Scientific, 34-1600), and VE-Cadherin (Santa Cruz Biotechnology, sc-6458) after fixation in cold methanol for 10 min followed by 3 PBS washes and 1 h incubation in TBS. Cells were maintained in a humidified 5% CO_2_ incubator at 37°C. Cells at low density were fed with fresh medium every other day; cells at high density were fed every day. Cells were split 1:2 or 1:3 at ≥80% confluency.

##### In vitro VCAM1 analysis

Bend.3 cells or primary BECs were seeded in 8-well chamber slides (154534, Thermo Scientific) overnight with 40,000 cells/cm^2^. The cells were serum starved for 1 h via incubation in DMEM with no added supplements (11965-092, Gibco), followed by treatment in DMEM with 10% pooled and dialyzed young (25 years or younger) or aged (65 years or older) human plasma (Bend.3 cells) or young (3-month-old) or aged (19-month-old) mouse plasma-derived serum (primary BECs and Bend.3 cells) for 16 h. Plasma was warmed to 37°C and filtered through 22 μm filter prior to being added to cells. The following day, cells were fixed in cold 4% PFA for 10 min, followed by three 5 min PBS washes and 1 h blocking in TBS with 3% donkey serum and .25% triton X-100 (TBS). Cells were blocked in TBS and primary antibody (1:250) overnight: anti-VCAM1 (ab19569, abcam) and anti-VE-Cadherin (sc-6458, Santa Cruz Biotechnology). Cells were blocked in TBS and secondary antibody (1:250) for 45 min the next day (Alexa Fluor 488: A10266, Life Technologies; Alexa Fluor 647: A10277, Life Technologies).

##### TNF-alpha dosage response

Bend.3 cells were seeded overnight as described above, serum starved for 1 h before being cultured for 24 h in varying concentrations of TNFα (5ng/mL-156.25 pg/mL) in DMEM, followed by staining with CD31 and VCAM1 antibodies and flow cytometry analysis.

##### In vitro flow cytometry

Bend.3 cells were serum starved for 1 h followed by 16 h overnight treatment with 10% young or aged mouse or human plasma-derived serum, as described above for in vitro VCAM1 analysis. After overnight treatment with plasma, medium was aspirated and cells were washed once with PBS. Cells were detached with 700 μl of accutase (A1110501, Life Technologies) for 5 min and the reaction was stopped by resuspending cells in 2 mL PBS. Cells were centrifuged for 5 min at 1100 rpm, medium was aspirated, and cells were resuspended in 1 mL/well of PBS with 4% PFA (diluted with 8 mL of PBS) and fixed on ice for 10 min. After centrifugation at 1100 rpm for 5 min, cells were resuspended in PBS for one wash followed by 30 min blocking in FACS buffer (PBS + 2% BSA with 2mM EDTA). Following centrifugation, cells were resuspended in 100 μl/sample of FACS buffer. FC blocking antibody (553142, BD Pharmigen) was added for 10 mins followed by addition of each antibody. Samples were incubated in antibodies for 30 min—1 h on ice. After two washes with FACS buffer, cells were resuspended in 500 μl FACS buffer and transferred to flow tubes for analysis. Bend.3 cells are stained using a conjugated anti-CD31 antibody (CD31-APC) (551262, BD Pharmigen), an anti-VCAM1 antibody (BE0027, BioXcell) conjugated using Dylight 488 Conjugation Kit according to manual instructions (53024, Thermo Scientific), and rat anti-mouse E-Selectin-DL405 (clone RB40.34, gift of the Butcher lab). Plasma treated Bend.3 cells were also stained with anti-CD31-PE/Cy7 (102418, Biolegend), anti-ICAM1-DL594 (clone YN1/1.7.4, gift of the Butcher lab), and anti-P-selectin-DL405 (clone 10E9.6, gift of the Butcher lab). Cells were gated on forward (FSC = size) and sideward scatter (SSC = internal structure). FSC-A and FSC-W blotting was used to discriminate single cells from cell doublets/aggregates. PI+ dead cells were excluded. CD31+ cells were defined as BECs.

#### Western blot analysis

Western blots are performed using cell lysate or whole tissue lysate from cortex and hippocampi. Cells are scraped in cold PBS and then spun down at 1100 rpm for 5 min. The cell pellet is then reconstituted in 100 μL of RIPA lysis buffer (89900, Pierce Biotechnology) with protease and phosphatase inhibitor (78441, Thermo Scientific) and 2 mM EDTA added and snap frozen to disrupt the cell membranes. Cells are then thawed on ice, and spun down at 10,000 rcf for 10 min. Supernatant is transferred to final tubes and mixed with 4X Staining Buffer and β-mercaptoethanol (1:5) after protein is saved for quantification. Protein quantification is performed using the Pierce BCA Protein Assay Kit and according to manual instructions (23225, Thermo Scientific). Protein lysate (approximately 18 μg) was run through 12- or 10-well 4–12% NuPage gels (NP0321BOX, Thermo Scientific) for approximately 1.5 h at 110 volts, followed by a semi-wet transfer onto nitrocellulose membrane (1620115, Bio-Rad) for 1 h and 15 min at 20 volts. Western blots were blocked in donkey anti-goat (sc-2020, Santa Cruz Biotechnology) and donkey anti-rabbit (sc-2780, Santa Cruz Biotechnology) secondary antibodies for in Tris-Buffered Saline with 0.05% Tween 20 (TBST) + 5% milk (1:10,000). Blots were imaged using Odyssey CLx and Image Studio software (Li-Cor). Pixel intensity quantification of western blots was done using ImageJ software.

#### Immunostaining

<u>Non-BrdU/EdU staining:</u> Brain sections were washed 3 times for 10 min in TBST and then blocked in TBS (TBS, 3% donkey serum – 130787, Jackson ImmunoResearch, 0.25% triton X-100 – T8787, Sigma Aldrich) for 1.5 h, followed by 72 h incubation on rocking platform at 4 °C in primary antibodies (see antibodies listed below). For secondary staining, brain sections washed 3 times for 10 min in TBST, followed by 1.5 h incubation in secondary antibody mix. Following secondary antibody incubation, there were 4 10 min washes in TBST; the second wash of which contained Hoechst (1:2000) with TBST. Brain sections were mounted on Superfrost™ microscope slides (12-550-15, Fisher Scientific) with Flouromount™ Aqueous Mounting Medium (Sigma-Aldrich, F4680). Slides were stored at 4°C.

<u>EdU staining:</u> Brain sections were washed 3 times for 10 min in TBST and then blocked in TBS (TBS, 3% donkey serum, 0.25% triton X-100) for 1.5 h, followed by 72 h incubation on rocking platform at 4 °C in primary antibodies (see antibodies listed below). For secondary staining, brain sections washed 3 times for 10 min in TBST, followed by 1.5 h incubation in secondary antibody mix. After secondary staining sections were washed 3 times in TBST followed by 50 min fixation with 4% PFA and 3 10 min washes in TBST. EdU staining was performed following manual instructions on the Imaging Kit (C10637, Life Technologies). 4 10 min washes, in which the third wash contained Hoechst (1:2000), with TBST before mounting with Flouromount™ and slides were stored at 4°C.

<u>BrdU staining:</u> Brain sections were washed 3 times for 10 min in TBST and then transferred to 95°C for 10 min in 10 mM sodium citrate (pH 6), followed by 3, 10 min washes in TBST. Sections were incubated in 3M HCl for 30 min at 37°C and then washed 3 times for 10 min washes in TBST. Sections were blocked for 1.5 h TBS and then transferred primary antibody mix for 72 h incubation on a rocking platform at 4 °C. Secondary staining started with 3 washes for 10 min in TBST, followed by incubation with secondary antibody mix for 1.5 h (all antibodies listed below). After 3, 10 min washes in TBST, sections were mounted with Flouromount™. Slides were stored at 4°C.

#### Enzyme-linked immunoabsorbent assay (ELISA)

Mouse plasma samples were used to measure soluble VCAM-1 using VCAM-1 ELISA kit (Raybiotech, ELM-VCAM-1) according to manual instructions. Soluble VCAM1 was measured in human plasma samples using the Human sVCAM-1/CD106 Quantikine ELISA Kit (R&D Systems, DVC00), following manual instructions. Optical density was measured at 450 nm and 540 nm wavelengths on a Varioskan Flash Multimode Reader (5250040, ThermoScientific).

#### Antibodies

**Primary:** Rat monoclonal anti-BrdU (1:500, Abcam, ab6326), Click-iT® Plus EdU Alexa Fluor® 488 Imaging Kit (Thermo/Life Technologies, C10637), goat monoclonal anti-Sox2 (1:100, Santa Cruz, sc17320), mouse monoclonal anti-GFAP (1:1000, Chemicon/Fisher, MAB360MI), rat monoclonal anti-VCAM1 (1:125, Abcam, ab19569), DyLight 488 Lectin (1:200, Vector, DL-1174), rabbit monoclonal anti-Aquaporin 4 (1:500, Millipore, AB2218), rat monoclonal anti-CD68 (1:600, Serotec, MCA1957), goat polyclonal anti-Iba1 (1:250, ProteinTech, 10904-1-AP), goat polyclonal anti-doublecortin (DCX) (1:100, Santa Cruz, sc8066), mouse anti-human-VCAM1 antibody (BBA5, Novus Biologicals), monoclonal mouse anti-human IgG antibody (MAB002, R&D Systems); rat monoclonal anti-VCAM-1 (clone M/K-2.7, Bioxell, BE0027); Rat IgG 1 Isotype antibody (Clone HRPN, Bioxell, BE0088), VE-Cadherin (sc-6458, Santa Cruz Biotechnology), FC blocking antibody (553142, BD Pharmigen), rat anti-CD31 antibody (CD31-APC) (551262, BD Pharmigen), Dylight 488 Conjugation Kit (53024, Thermo Scientific), Anti-Mouse CD45 PerCP-Cyanine5.5 (1:1000, eBioscience, 45-0451-80), PerCP/Cy5.5 anti-mouse CD11a/CD18 (LFA-1) (1:100, Biolegend, 141007), Anti-Mouse CD11b PerCP-Cyanine5.5 (1:100, eBioscience, 45-0112-80), Anti-Mouse TER-119 PerCP-Cyanine5.5 (1:100, eBioscience, 45-5921-80), CD13 Antibody (ER-BMDM1) APC (1:50, NOVUS Biologicals, NB100-64843), Anti-ACSA-2-PE mouse (clone: IH3-18A3) (1:100, Miltenyi Biotec Inc., 130-102-365), Anti-Mouse CD31 (PECAM-1) PE-Cyanine7 (1:100, eBioscience, 25-0311-81), Anti-mouse MECA-99 antibody was a gift of the Butcher lab and labeled with fluorophores using DyLight™ Antibody Labeling Kit (DyLight™488, Thermo Scientific, 53025), CD31-APC (1:100, BD 551262), CD45-FITC (1:100, BD Pharmingen Clone 30-F11 553080), and Cd11b-BV421 (1:100, Biolegend Clone M1/70 101236)

**Secondary:** Alexa Fluor® 488 donkey anti-goat IgG (1:250, Thermo/Life Technologies, A-11055), Alexa Fluor® 488 donkey anti-rat IgG (Invitrogen/Life Technologies, A21208), Alexa Fluor® 555 donkey anti-mouse IgG (1:250, Invitrogen, A31570), Alexa Fluor® 555 donkey anti-goat IgG (1:250, Invitrogen, A21432), Cy3 AffiniPure donkey anti-rat IgG (1:250, Jackson Immunoresearch, 712-165-153), Alexa Fluor® 647 donkey anti-mouse IgG (1:250, Invitrogen, A31571), Alexa Fluor® 647 donkey anti-rabbit IgG (1:250, Life Technologies, A-31573), Alexa Fluor® 647 donkey anti-goat IgG (1:250, Invitrogen, A-21447), Alexa Fluor® 647 donkey anti-rabbit IgG (1:250, Life Technologies, A-31573), Alexa Fluor® 647 donkey anti-goat IgG (1:250, Invitrogen, A-21447), Alexa Fluor 488 Azide (A10266, Life Technologies); Alexa Fluor 647 Azide (A10277, Life Technologies); Hoechst 33342 (1:2000, Sigma, 14533-100MG)

##### Quantitative analysis of immunohistological data

#### Neurogenesis

For quantification of the number of BrdU+, EdU+, Sox2+, DCX+, NeuN+, and GFAP+ cells in mice, confocal Z-stacks of six coronal brain sections spanning the hippocampus (40 mm thick, 200 mm apart) were captured on a Zeiss confocal microscope, and cells within the dentate granule cell layer were counted. Neurogenesis was quantified by counting colabeled precursor cell populations in the neurogenic granular layer and hilus in 6 serial 40 μm sections throughout the hippocampal DG. Counts were made while blinded to the different groups. Cell numbers were normalized to the volume of the DG granule cell layer. Unless otherwise noted, all representative confocal images were captured on a Zeiss confocal microscope and represent a Z-stack of 10 images taken on confocal planes 2 μm apart. Images of cells cultured in vitro (Supplemental Figure 4) were captured on a single plane.

**For 3D Rendering of High-magnification Immunofluorescence Tissue Staining (Supplemental Figure 6):** 63x magnification Z-stacked Images were captured using the LSM 700 system set to take 51 slices with an interval of 0.4 um. 3D modeling through the Imaris8 software was done on the Z-stacked Images for high contour modeling of antibody markers. Surface masks with smoothing set to 0.2 um were used to form the base 3D model. Surface area and voxel thresholds were used to remove artifacts from high magnification image acquisition.

For %area quantifications of immunofluorescent images, ImageJ was used to determine integrated pixel intensity of thresholded images comprising 5–6, serial 40 μm sections spanning the hippocampal DG per animal immunostained for various vasculature and microglia cell markers.

#### Vasculature

VCAM1+ and Lectin+ signals were threshold and analyzed using the “% Area” of the ‘Analyze Particles’ function in ImageJ 1.50i (ImageJ website: https://imagej.nih.gov/ij/). Auto-fluorescent background picked up by the thresholds is removed using the Size Exclusion feature of the ‘Analyze Particles’ function. Colocalized VCAM1+/Lectin+ signal was calculated using the Colocalization plug-in developed by Pierre Bourdoncle of the Institut Jacques Monod (Plug-in website: http://rsb.info.nih.gov/ij/plugins/colocalization.html).

#### Microglial Activation

Microglia markers IBA1 and CD68 were analyzed through “% Area” and manual cell counts; IBA1+ and CD68+ signals were threshold and analyzed using the “% Area” of the ‘Analyze Particles’ function of ImageJ 1.50i (ImageJ website: https://imagej.nih.gov/ij/). Colocalized IBA1+/CD68+ signal was calculated using the Colocalization plug-in (Plug-in website: http://rsb.info.nih.gov/ij/plugins/colocalization.html). To count the numbers of IBA1+ and IBA1+/CD68+ cells in the DG, the soma of IBA1+ cells in 5–6 visual fields per animal photographed using a Zeiss confocal microscope and Zen Black software were physically counted using the ‘Multi-point’ tool of ImageJ 1.50i (ImageJ website: https://imagej.nih.gov/ij/). Colocalization of IBA1+/CD68+ cells was counted through visual confirmation by superimposition of the two markers in ImageJ. Cell density of IBA1+ and IBA1+/CD68+ cells in the DG were calculated by normalizing the total counts to the volume of the DG determined by the dimensions of the images captured using the Zeiss confocal microscope.

Data were analyzed using an unpaired Student’s t-test and One- or Two-Way Anova with Tukey’s Multiple Comparisons Test or Sidak’s Multiple Comparisons Test, respectively. P values equal to or lower than .05 were considered statistically significant. Significance was assessed based on p values and heteroscedastic variance between groups that are statistically compared.

